# Genome-Wide Association Study of Brain Connectivity Changes for Alzheimer’s Disease

**DOI:** 10.1101/342436

**Authors:** Samar S. M. Elsheikh, Emile R. Chimusa, Nicola J. Mulder, Alessandro Crimi

## Abstract

Variations in the human genome have been found to be an essential factor that affects susceptibility to Alzheimer’s disease. Genome-wide association studies (GWAS) have identified genetic loci that significantly contribute to the risk of Alzheimers. The availability of genetic data, coupled with brain imaging technologies have opened the door for further discoveries, by using data integration methodologies and new study designs. Although methods have been proposed for integrating image characteristics and genetic information for studying Alzheimers, the measurement of disease is often taken at a single time point, therefore, not allowing the disease progression to be taken into consideration. In longitudinal settings, we analyzed neuroimaging and single nucleotide polymorphism datasets obtained from the Alzheimer’s Disease Neuroimaging Initiative for three clinical stages of the disease, including healthy control, early mild cognitive impairment and Alzheimer’s disease subjects. We conducted a GWAS regressing the absolute change of global connectivity metrics on the genetic variants, and used the GWAS summary statistics to compute the gene and pathway scores. We observed significant associations between the change in structural brain connectivity defined by tractography and genes, which have previously been reported to biologically manipulate the risk and progression of certain neurodegenerative disorders, including Alzheimer’s disease.

## Introduction

Alzheimer’s disease (AD) is a neurodegenerative disease with believed onset in the hippocampus. It subsequently spreads to the temporal, parietal, and prefrontal cortex^1^. Symptoms of the disease worsen over time, and as the patient’s condition declines, AD ultimately leads to death. Causes of the disease are yet unclear, and it has even been hypothesised to be related to external bacteria^2^. However, 70% of AD risk is believed to be contributed by complex genetic risk factors^3^. The protein encoded by the apolipoprotein E (*APOE*) gene, located on chromosome 19, carries cholesterol in the brain, affecting diverse cellular processes. Carriers of the *APOE* allele *ε*4 have three times the risk of developing AD compared to non-carriers^4^. Although *APOE ε*4 is the primary genetic risk factor that contributes to the development of late-onset AD; its effect accounts for only 27.3% of the overall disease heritability, which is estimated to be 80%^5^.

In order to estimate the remaining heritability of AD, many attempts have been made to uncover additional genetic risk factors. Genome-wide studies have successfully identified single nucleotide polymorphisms (SNPs) which affect the development of AD^6–8^. Understanding the underlying biological process of the disease, and identifying more potential genetic risk variants, could contribute to the development of disease-modifying therapies.

On the other hand, the recent advancements in imaging technologies have provided more opportunities for understanding the complexity of how the brain connects, and at the same time, enhancing and forming a more reliable basis for neuroimaging and human brain research^9^. By merging brain imaging with genetics, previous studies proposed different ways of analyzing the data, to discover genetic factors that affect the structure and function of the human brain. Significant efforts in this area have been made by the Enhancing Neuroimaging Genetics through Meta-Analysis (ENIGMA)^10^ project. The methods offered several diverse ways to link together two heterogeneous collections of data - brain imaging and genetic information - depending on the hypothesis under study, and hence, the type of images and genetic information.

Stein et al.^11^ used the T1 weighted Magnetic Resonance Imaging (MRI) scans from the Alzheimer’s Disease Neuroimaging Initiative (ADNI)^12^ and developed a voxel-based GWAS method (vGWAS) that tests the association of each location in the brain (each voxel), with each SNP. To quantify the phenotype, they used the relative volume difference to a mean template at each voxel, and their method, vGWAS, can be applied to other brain maps with coordinate systems. Although vGWAS did not identify SNPs using a false discovery rate of 0.05, they highlighted some genes for further investigation. More recently, other studies on the genetics of brain structure implemented a genome-wide association of the volume in some sub-cortical regions, and successfully identified significant genetic variants^13, 14^. Additional efforts in the literature include the development of multivariate methods that aim at identifying the imaging-genetics associations through applying sparse canonical correlations, to adjust for similarity patterns between and within different clinical stages of the disease - an assumption which is hardly met^15^.

Connectomics^16^, or the study of the brain connectome, is a novel advancement in the field of neuroimaging. A structural or functional brain connectome is a representation of the brain, and its connections, as one network. The connectome comprises nodes representing different and distinct regions in the brain, and edges representing the functional or structural connection between brain regions. More specifically, the edges of a structural connectivity network are defined by the anatomical tracts connecting the brain regions^17^. Those are extracted from diffusion weighted imaging (DWI), a type of imaging which detects the diffusion of the water molecules in the brain. Furthermore, the connectome can be summarised by several global and local network metrics^17^ which allow the study of the brain as one entity (one scalar value), the comparison of different groups of participants, as well as the study of variation between and within different brain regions. DWI is not the only method to represent structural connectivity. Structural connectivity can be defined by using T1 scans and voxel-based morphometry (VBM)^18^, a technique investigating structural tissue concentration, especially in gray matter (GM). It has been demonstrated that the morphology across the brain is governed by covariation of gray matter density among different regions^19^. In this way a structural connectome is constructed defining the edges among brain regions as the correlation of GM morphology. This can complement the DWI approach which is mostly based on white matter. Of particular interest are covariability hubs, nodes of this network which have high degree centrality since they are the most representative of the overall cortex^19, 20^.

In an attempt to understand aging of the healthy brain, Wu et al.^21^ carried out a longitudinal study of the structural connectome in healthy participants. Their analysis evaluated the association of the annual change in both the local and global network characteristics with age, but no genetic investigation was carried out in relation to those longitudinal features. In another study Jahanshad et al.^22^ conducted a GWAS on dementia subjects using connectivity patterns as a phenotype, and identified the genetic variant rs2618516 located in the *SPON1* gene; however, this study considered cross sectional phenotype, collected at one specific time point. VBM-based GWAS have also been carried out comparing AD and control subjects at one specific time point^23^, identifying the *APOE* gene and other SNPs related to ephrin receptor as markers strongly associated with multiple brain regions.

In this paper, we used a dataset from ADNI (http://adni.loni.usc.edu/) to perform four quantitative GWAS, with the longitudinal change in the brain connectome used as a phenotype. Our choice of using the ADNI dataset was because the particular combination of data types needed to run this analysis was available, in the context of AD. We used the absolute difference in the longitudinal integration and segregation global network metrics to represent the change in structural brain connectivity defined by tractography. After obtaining the GWAS summary statistics for all the SNPs typed in the original data, we aggregated their p-values using the PAthway SCoring ALgorithm (PASCAL) software^24^ and computed genome-wide gene and pathway-scores. Our result identified a number of genes significantly associated with the change in structural brain connectivity, including *ANTXR2*, *OR5L1*, *IGF1*, *ZDHHC12*, *ENDOG* and *JAK1*. Most of those genes were previously reported to biologically manipulate the risk and progression of certain neurodegenerative disorders, including Alzheimer’s disease^25–28^. Additionally, we investigated whether there are additional changes in connectivity defined by GM covariability.

## Results

### Analysis Pipelines

In this work, we used a longitudinal imaging dataset, combined with genetic variation information at the SNP-level. The sample consists of three groups which represent three distinct clinical stages of Alzheimer’s disease. This includes healthy individuals (controls), Early mild cognitive impairment (MCI), and Alzheimer’s disease. Aiming at studying the genetic effect on the longitudinal change in the brain structure for those groups, we conducted genome-wide tests of the associations between brain image features and different levels of genetic variations. These image features were derived from an intensive map of the brain’s neural connections. The overall pipeline followed is summarized in Fig. 1.

**Figure 1.**
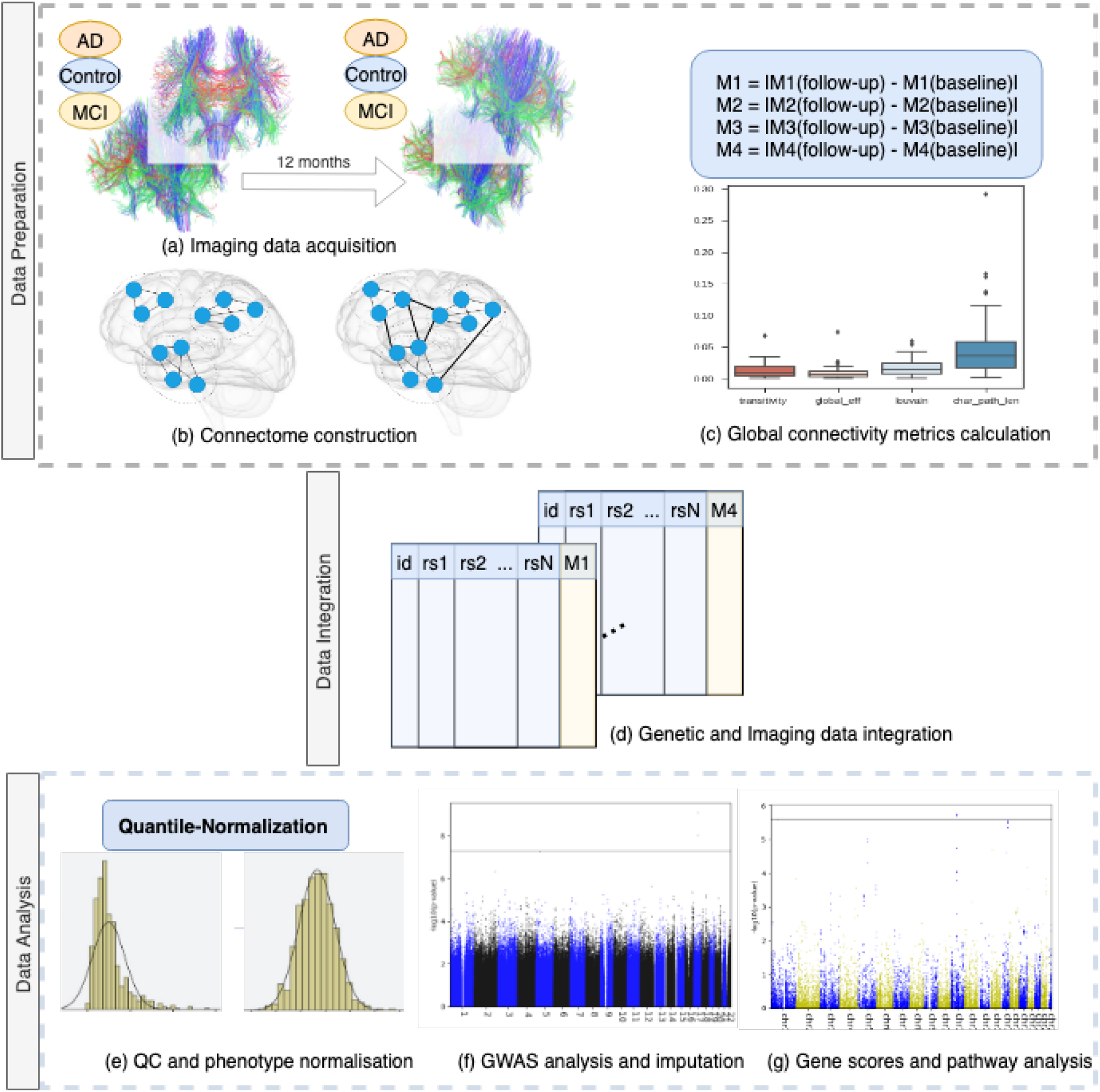
The analysis pipeline: (a) The DWI images were collected at two time points, for three clinical stages of AD. (b) The images were processed using distinct brain regions from the Automated Anatomical Labeling (AAL) atlas, and two structural connectomes were constructed for each participant at each time point. (c) Global connectivity metrics were computed, along with the absolute difference between the baseline and follow-up measures. (d) The latter were merged (as phenotypes) with the PLINK FAM files for all subjects present in both datasets. (e) All essential quality control procedures were performed before GWAS analysis, besides the quantile normalization of phenotypes. (f) GWAS was conducted using PLINK, and, (g) the resulting summary statistics were used by PASCAL software to calculate the gene- and pathway-scores accounting for LD patterns using a reference dataset.

### Descriptive Statistics of Brain Imaging Features

Using the DWI images at both baseline and a follow-up visit after 12 months, the brain connectome was constructed. We obtained four global network metrics, as explained in the Methods section. We chose network transitivity and Louvain modularity to represent network segregation, along with characteristic path length and weighted global efficiency to represent the brain integration^17^.

Each of these four metrics quantitatively represents the whole brain network as a single value. Supplementary Fig. **??** illustrates both the distribution of the network metrics in the data, in the baseline and follow-up, for the three participant categories. The figure also shows the association patterns of the metrics. A similar figure that illustrates the distribution of the absolute difference between the baseline and follow-up metrics is shown in Supplementary Fig. **??**, it compares the three groups in each sub-figure. To determine how the differences between these connectivity metrics are distributed, we plotted four boxplots, as shown in Supplementary Fig. **??**, for the three groups combined.

To verify that the longitudinal change is consistent and not the result of artifacts, we initially compared the imaging features between the two-time points. Table 1 shows the results of the non-parametric Wilcoxon test between baseline and follow-up features. The test ranks the values of the paired measurements and compares their central values. In this test, we used the AD patients and controls, most of the metrics turned out to have significant longitudinal differences in the AD brain, but not in a healthy brain. Figure 2 compares the four metrics at the two time points, for each group individually, utilizing their boxplots.

**Table 1.** Non-parametric Wilcoxon test of the difference between brain connectivity features at baseline and follow-up

| Group | P-values with * are significant (< 0.05) |  |  |
| --- | --- | --- | --- |
|  | Network Metric | Statistic | P-value |
| **AD** | Characteristic path length | 107.0 | 0.0057* |
|  | global efficiency | 98.0 | 0.0033* |
|  | Transitivity | 226.0 | 0.6664 |
|  | Louvain | 114.0 | 0.0086* |
| **MCI** | Characteristic path length | 612.5 | 0.0891 |
|  | global efficiency | 672.0 | 0.2196 |
|  | Transitivity | 760.0 | 0.5972 |
|  | Louvain | 712.0 | 0.3630 |
| **Control** | Characteristic path length | 496.0 | 0.2465 |
|  | global efficiency | 55.0 | 0.1172 |
|  | Transitivity | 517.0 | 0.3421 |
|  | Louvain | 529.0 | 0.4062 |

**Figure 2.**
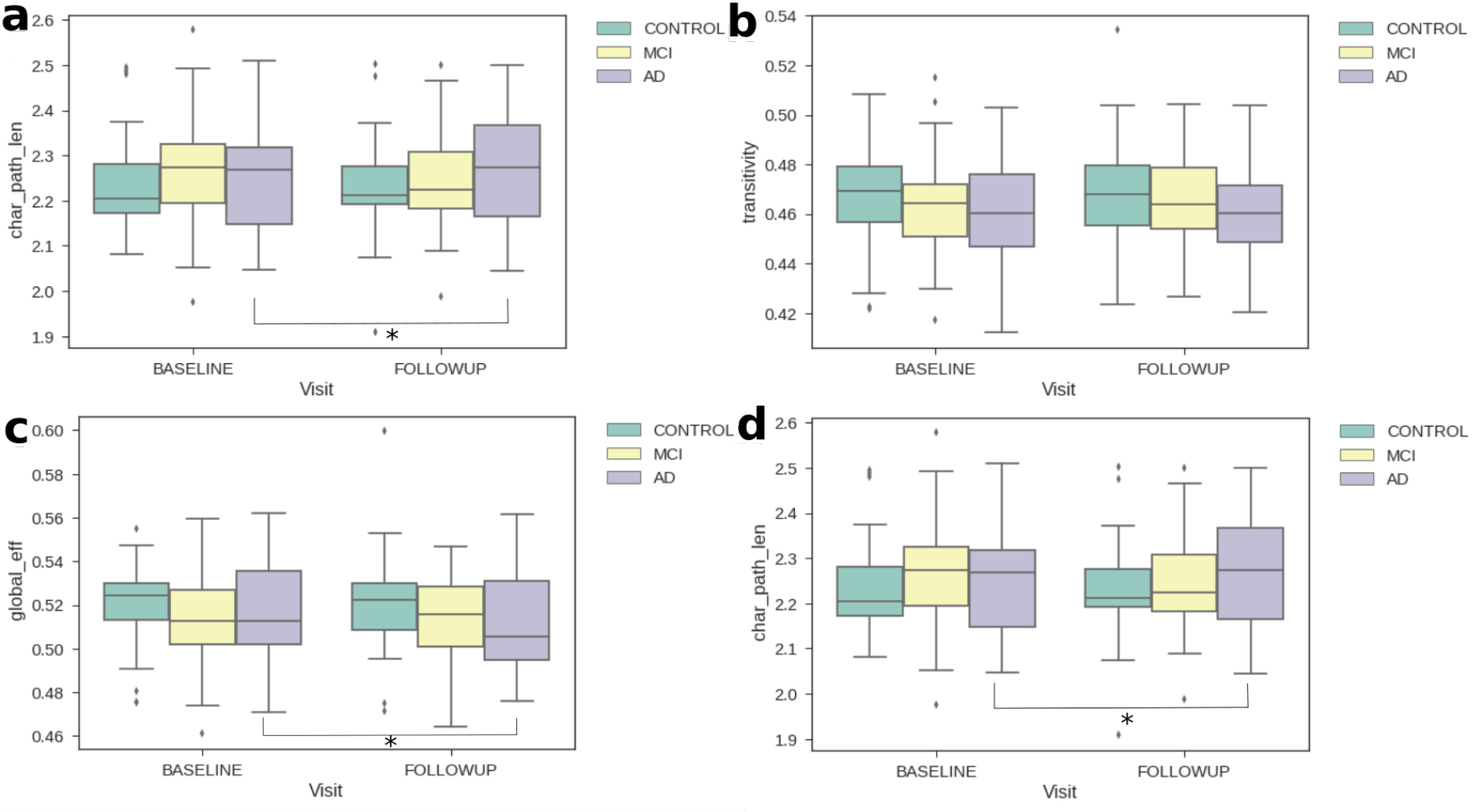
Boxplots for global network metrics to compare AD and controls in the baseline (green) and follow-up (yellow). The metrics are, Louvain modularity (a), transitivity (b), global efficiency (c) and characteristic path length (d). It is evident that at least the means for the AD population are different while for the others they are generally unvaried. The asterisk denotes that there is a significant change from baseline to the follow-up visit (p-value< 0.05).

We further investigated structural connectivity given by VBM, namely whether there is a structural covariability change in the gray matter. Before doing this, a traditional VBM analysis was carried out. In particular, we compared the AD against the control population at the two time points, and the two populations individually compared to themselves at the two time points. The differences between the groups were given by a t-test converted into corrected p-values^29^. Between the same populations at different time points no significant voxels were found, but comparing the two different populations, most of the brain regions were statistically significantly different. Figure 8 show the t-statistics map of these differences. In line with previous work^30^, we identified the peak of statistical difference between AD and control subjects in the hippocampus, followed by the cingulate cortex and the temporal lobe at both time points. The hubs detection was conducted on the same segmented GM data used in the VBM analysis, and again no significant differences were noted within the same population comparing different time points. The average hubs index is reported in Figure 9, and supplementary Figure **??** depicts the values averaged according to the ROIs of the AAL atlas, specifically showing the hubs index for the AD population at baseline and followup (similar results were obtained for the control population). Here, the highest values, in line with similar results of previous studies on healthy volunteers, were in the fronto-lateral cortex, cingolum,^20^ and basal ganglia^19^. Given the fact that no statistical difference was found for longitudinal changes, both using the traditional VBM analysis and the cortical hubs, no feature of this kind was available for the integrated analysis, which was therefore focused on the structural connectivity given by the tractography.

### Integrated Analysis

After we obtained our phenotypes of interest, given by the longitudinal changes of the features between the two time points, we prepared our data for genome-wide association analysis (see Fig. 1) by first integrating the phenotypes and genotypes.

The necessary quality control procedures that precede GWAS analysis were run as explained in the Methods section. Briefly, they include cleaning the data such as removing all SNPs with small sample sizes, and individuals with relatedness, as well as population stratification correction. Fig. 3 shows the plots after correcting for the population stratification. We quantile-normalised our phenotypes to allow the use of the linear model in GWAS.

**Figure 3.**
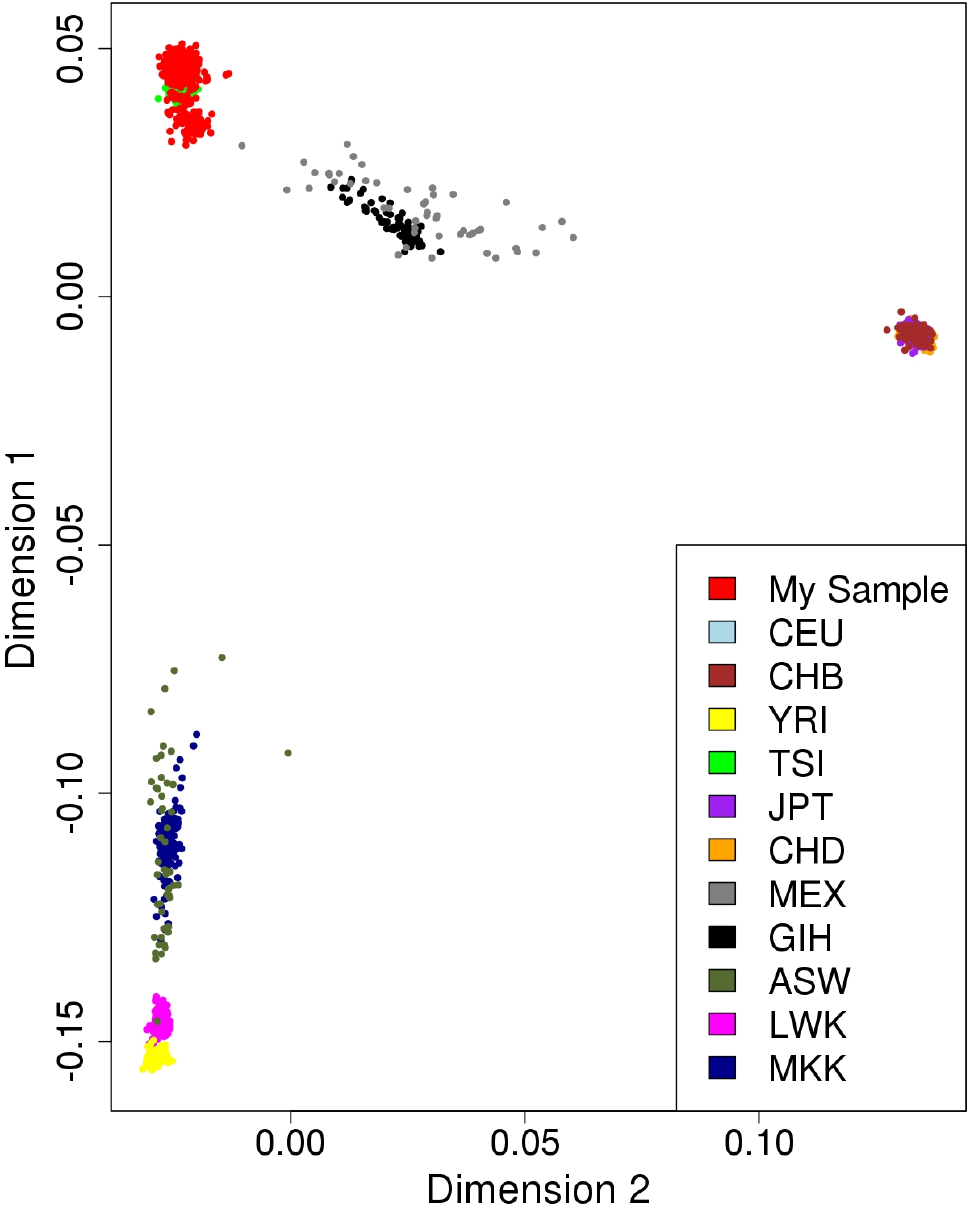
Quality control procedures: The plot shows the estimated ancestry of the genotypes of each study sample (in red) after applying the Multi-Dimensional Scaling (MDS). It also compares the genotype of the samples with a multiple ancestry reference. We observed that most of our participants belong to the Caucasian population, denoted here as CEU. A description of the reference population is found in the *Quality Control Correcting for Population Stratification* sub-section.

#### Genome-Wide Association Analysis

GWAS was conducted by regressing the normalised longitudinal changes of global connectivity metrics (response variable) on the SNPs’ minor allele frequencies (independent variable), one SNP at a time. Using PLINK^31^ we conducted four quantitative GWAS - one for each network metric, after which we performed a Gaussian imputation of GWAS summary results. Figure 4 shows the imputed GWAS results for the change in brain segregation metrics. The Manhattan sub-plots appear on the left, while the corresponding quantile-quantile (qq)-plots are on the right. Figure 5 shows the imputed GWAS results for brain integration metrics. The x-axis of the Manhattan plot represents the physical location along the genome, while the y-axis is the (−log 10(*p-value*)), and each dot represents a single SNP. In the qq-plots, the diagonal line represents the expected (under the null hypothesis) distribution of p-values, and similar to the Manhattan plot, each dot in the qq-plot represents a single SNP.

**Figure 4.**
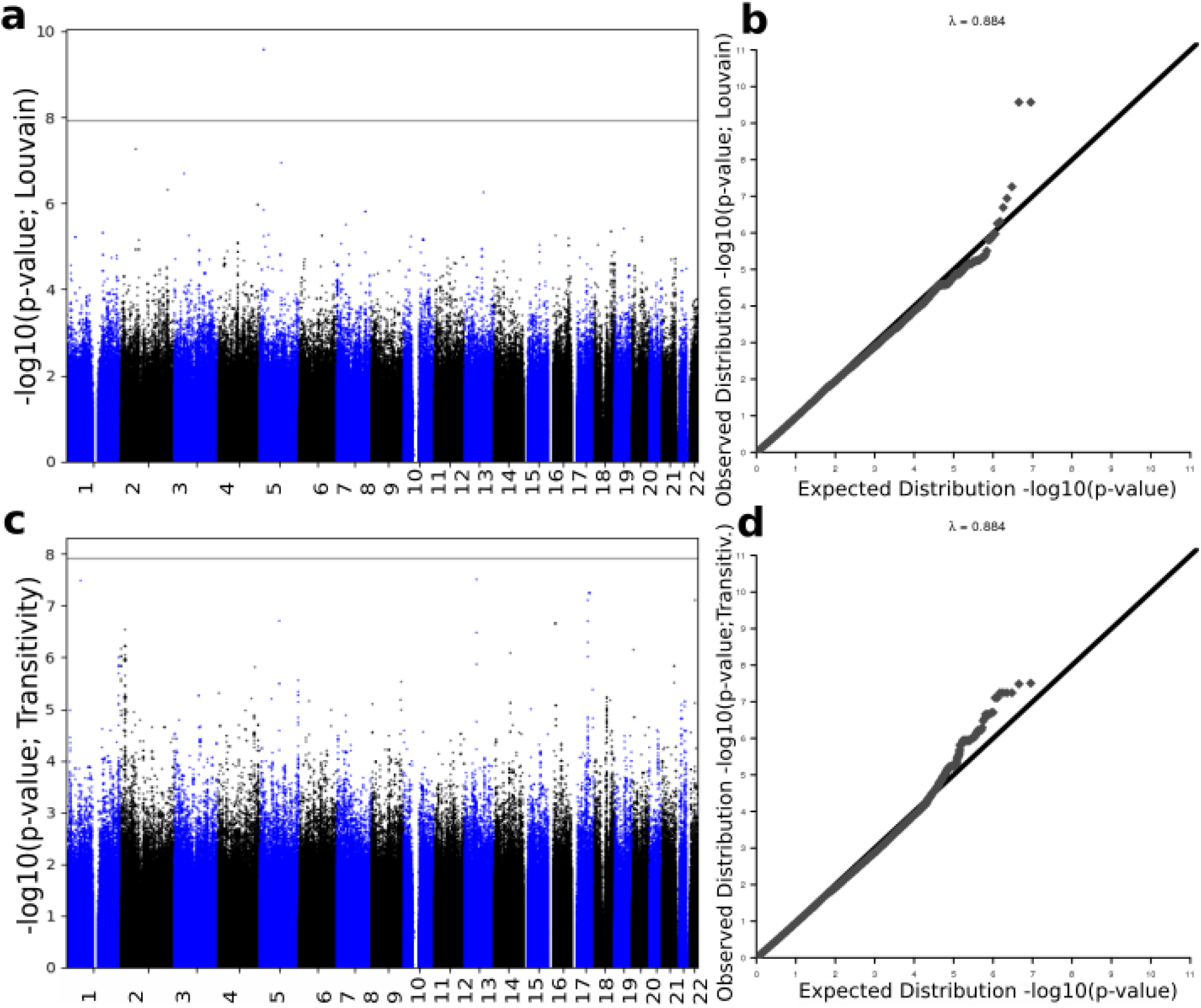
Imputation results of GWAS summary statistics for the change in segregation metrics. Top plots represent the change in Louvain modularity phenotype Manhattan plot (a and c) and quantile-quantile (qq)-plot (b and d). Bottom plots represents the change in transitivity phenotype. Louvain modularity imputation results show small evidence of deviation of measures before the tail of the distribution.

**Figure 5.**
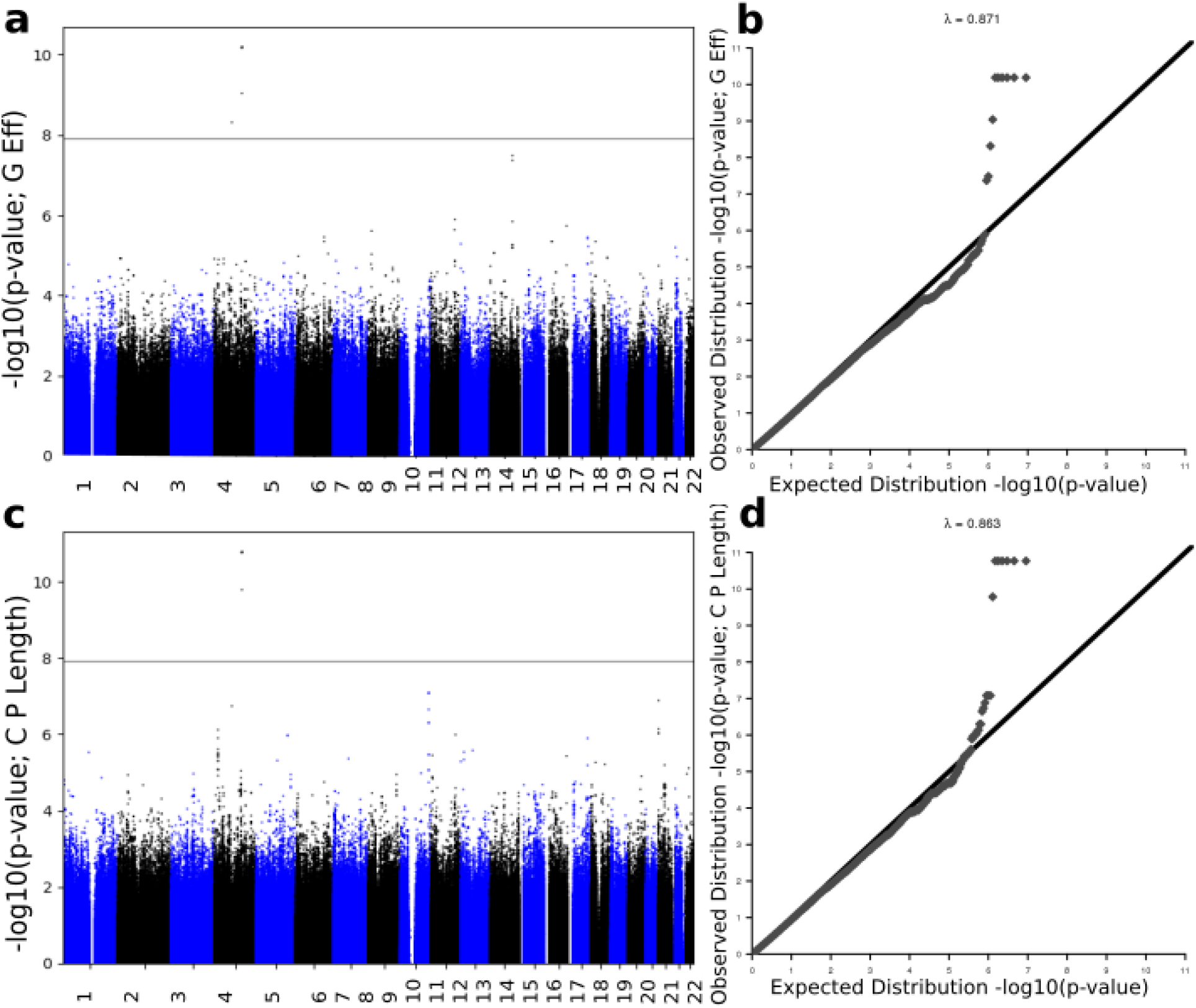
Imputation results of GWAS summary statistics for the change in integration phenotypes. Top plots represent the change in global efficiency Manhattan plot (a and c) and qq-plot (b and d), while the plots at the bottom represent the change in characteristic path length phenotype. Both qq-plots show very little evidence of deviation before the tail of the distribution.

The top 15 SNPs, including the significantly associated SNPs obtained after imputation of GWAS p-values for the absolute difference in Louvain modularity, transitivity, global efficiency and characteristic path length, are shown in Supplementary Tables **??**, **??**, **??** and **??**, respectively. The actual GWAS Manhattan plot for the absolute difference in segregation and integration metrics before imputation is provided in Supplementary Fig. **??**.

#### Gene and Pathway Scores

Using the imputed GWAS association results (p-values), we computed genome-wide gene scores, along with the pathway (gene set) scores, using the PASCAL software^24^. Figure 6 and Fig. 7 show the gene scores obtained for brain segregation and integration phenotypes, respectively.

**Figure 6.**
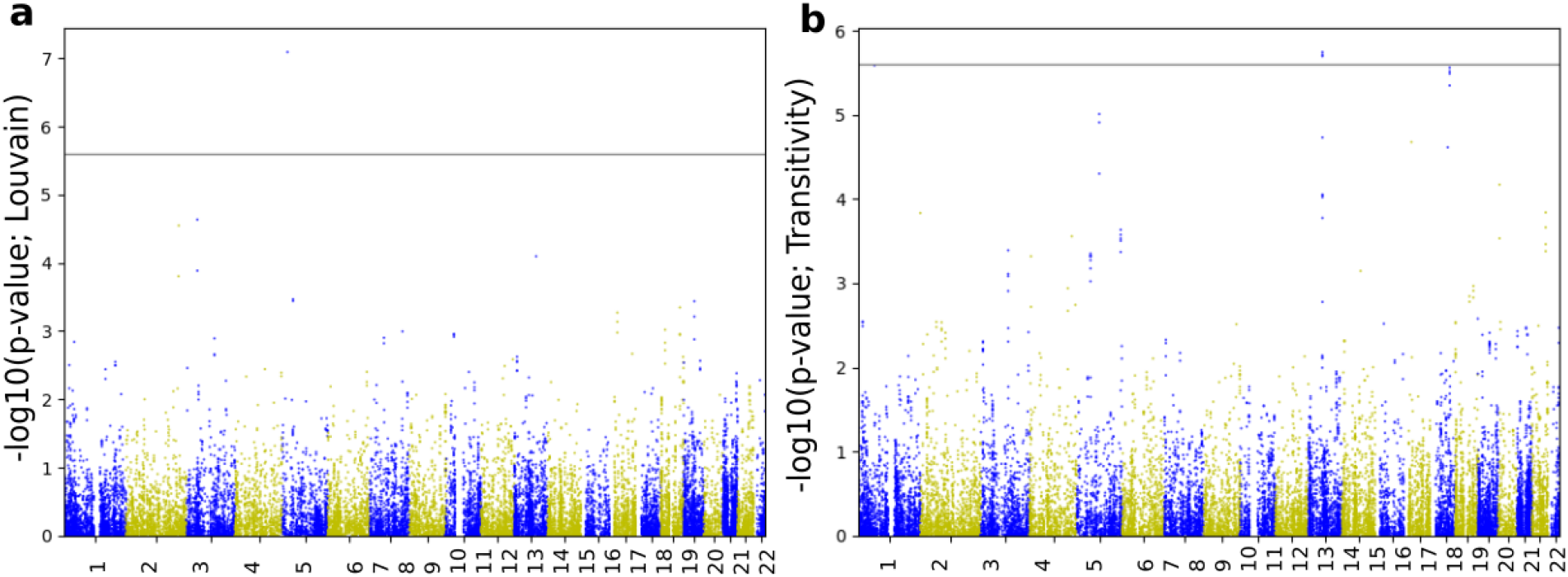
Manhattan plots of gene scores derived from imputed summary statistics for the change in segregation metrics. Lovain modularity appears in plot (a), and transitivity is illustrated by plot(b). The horizontal line represents the statistical threshold used here (2.5*E −* 6).

**Figure 7.**
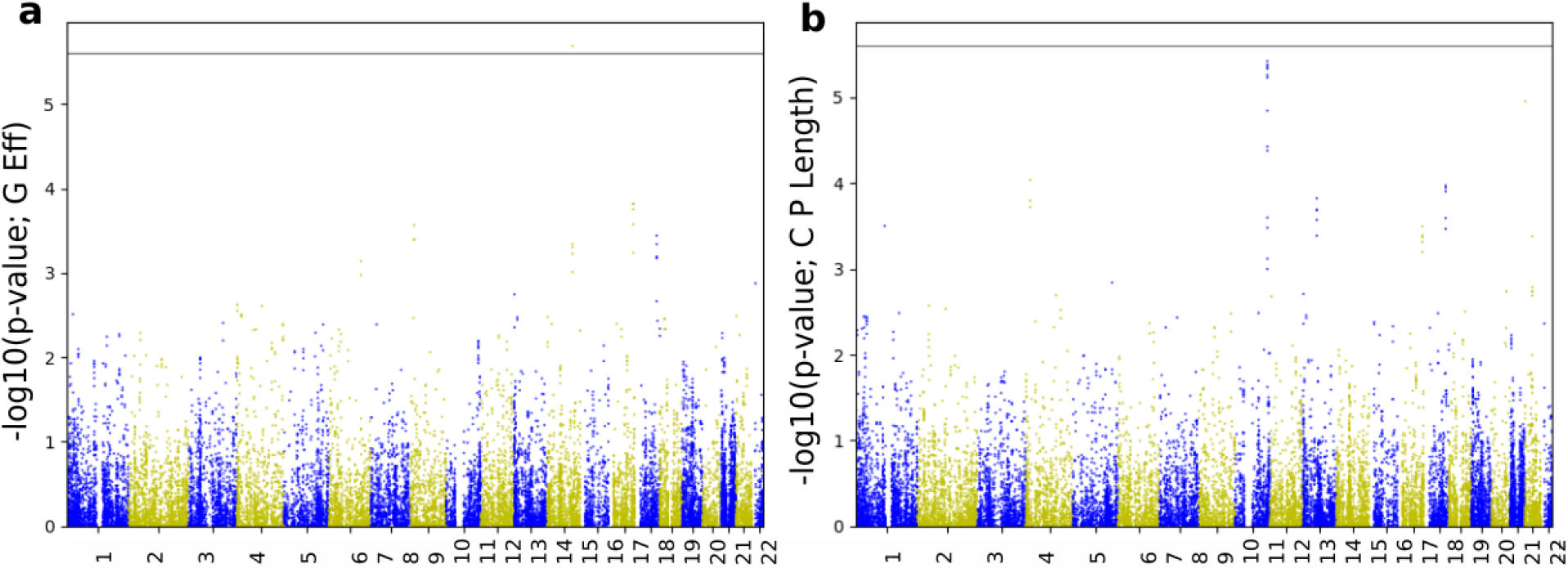
Manhattan plots of gene scores derived from imputed summary statistics for the change in integration metrics. Global efficiency is shown in plot (a), and characteristic path length is illustrated by plot (b). The horizontal line represents the statistical threshold used here (2.5*E −* 6).

**Figure 8.**
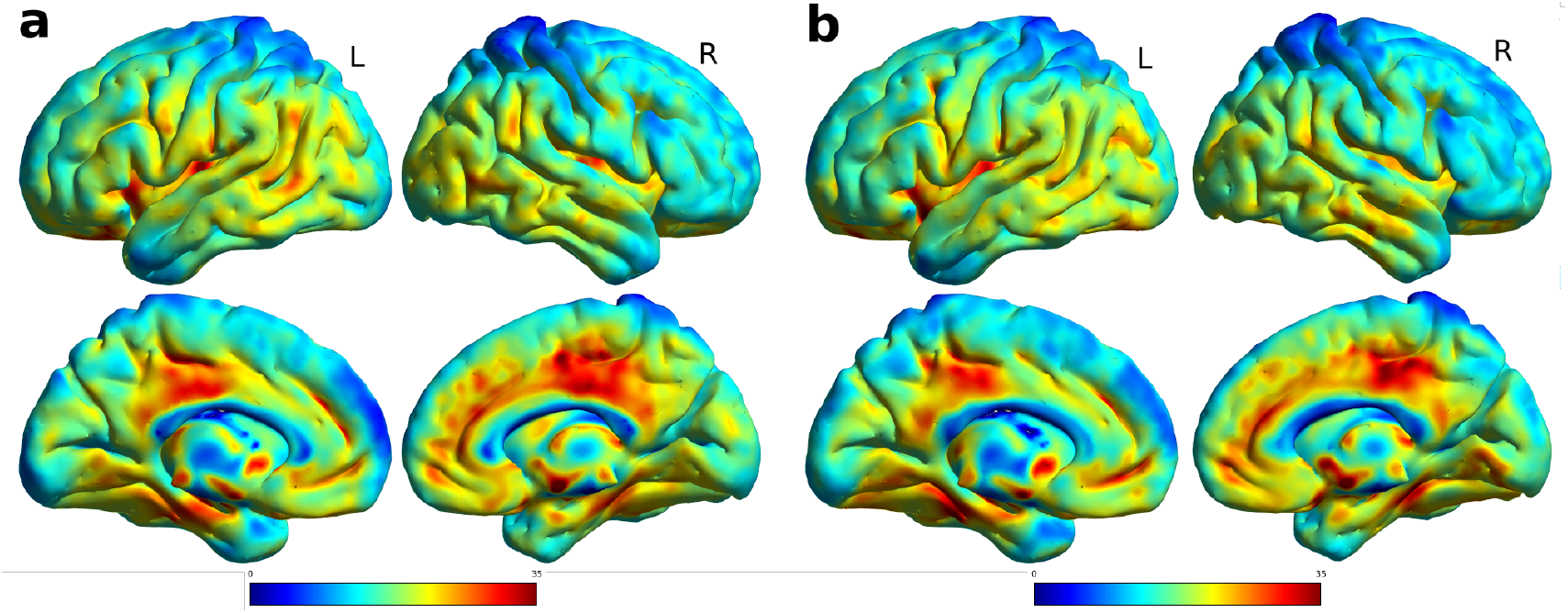
T-statistics map of the comparison between the VBM features of AD and control subjects. On the left (a) is the comparison at baseline, and (b) on the right for the followup. All views are for both hemispheres, lateral and medial view. Highest values, depicted in red, were at the hippocampus, cingulate cortex and temporal lobe for both time points.

**Figure 9.**
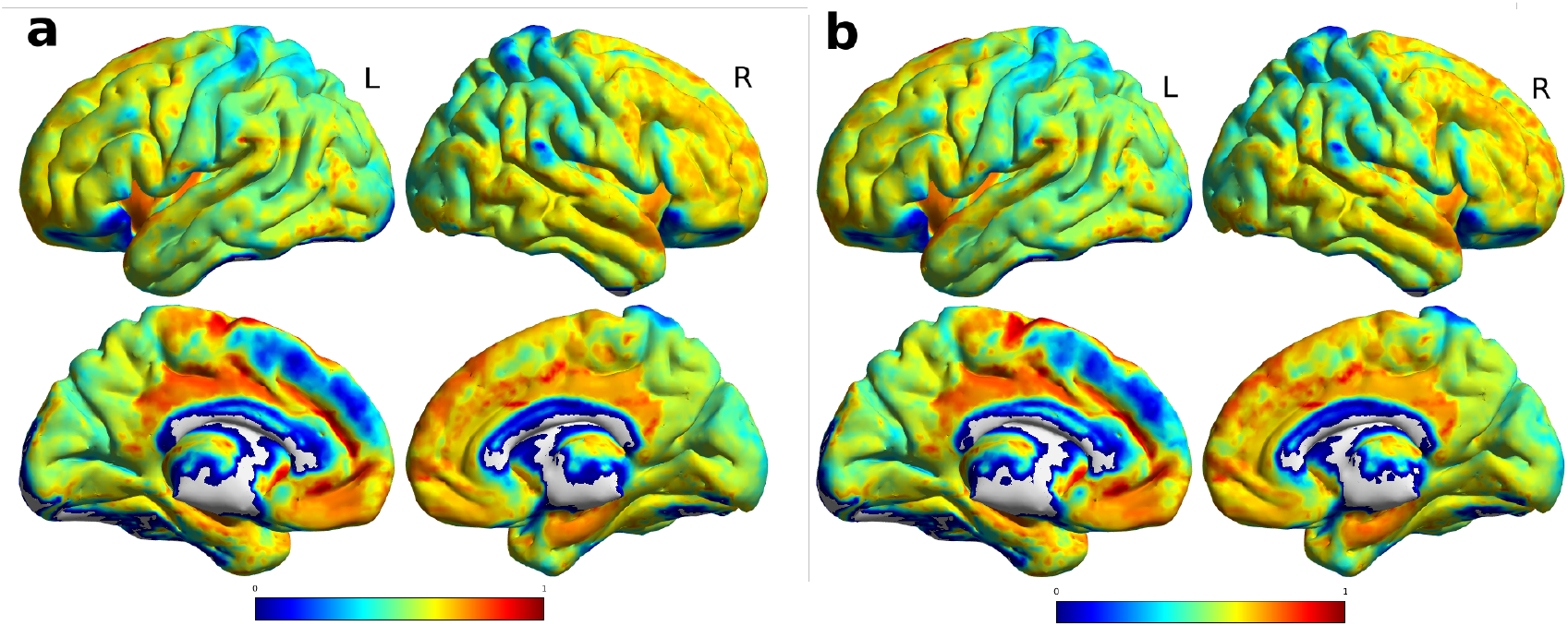
Average normalized connectivity hubs, (a) on the left there is the average value at baseline, and (b) on the right for the followup. All views are for both hemispheres, lateral and medial view. Highest values, depicted in red, were at the cingulate cortex, fronto-lateral cortex and basal ganglia, gray areas depict values of 0. The individual values averaged according to the ROIs of the AAL atlas are reported int the supplementary Figure **??**.

Using the total number of genes in the human genome (20, 000) we calculated the threshold. Therefore, we obtain the 5% gene-wide significance threshold by dividing the significance level by the total number of genes (or, tests), i.e.,

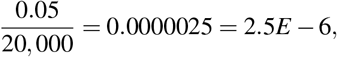

If we consider less power (90%) and 10% significance level, we get a gene-wide threshold of

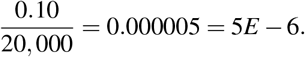

For each gene score result (and for both brain segregation and integration measures) we sorted our results and constructed a table of the top 30 genes (Table 3). The table also shows the 5% and 10% significant genes.

In the pathway results obtained for each metric and each chromosome, the total number of pathways used at each step was 1078. Therefore, the 5% threshold is 0.000046382 =4.6*e*− 5, while the 10% threshold is 0.000092764 = 9.28*e*−5. The gene *CDH18*, contains 3974 SNPs as per the data, was significantly associated with Louvain modularity change over time (p-value ≈ 8.09). On chromosome 11 and chromosome 15, a number of genes were associated with the change in brain connectivity through transitivity, while chromosome 9 shows a number of significant association results with characteristic path length. Supplementary Table **??** reports all the significant results as well as the top 20 pathways along the whole genome and in all the four phenotypes. As shown in the table, REACTOME BIOLOGICAL OXIDATIONS pathway, which consists of genes involved in biological oxidationsat was significantly associated with the change in Louvain modularity at 5% significance level (p-value=2.91E-5), on chromosome 10.

## Discussion

Association studies of human genome variation and imaging features of the brain have led to new discoveries in AD disease susceptibility. Previous GWAS and Next Generation Sequencing (NGS) identified about 20 genetic loci risk factors associated with AD^32^. More recently, cross-sectional studies of GWAS of the brain connectome successfully identified correlations between genetic variants and both AD and dementia^22^. Incorporating imaging features in a longitudinal setting with genetic information facilitates the identification of additional genetic risk factors which affect AD progression^33^. Here, we aim to identify the genetic variations which associate with AD brain neurodegeneration over time. The latter is measured as the change in global network metrics of the brain connectome of three clinical stages of AD.

In this study, we examine the significance of the change in the global network metrics over time, through Wilcoxon test statistics (shown in Table 1). We tested the distribution of each metric before and after one year, and only the AD brain showed a difference, compared to controls. We proceeded with the analysis by conducting four quantitative genome-wide association tests, taking the absolute difference in the metrics of brain network integration and segregation as individual phenotypes. To our knowledge, this is the first study of its type, to compare longitudinal imaging features of the connectome to genetic information. These connectivity features were obtained from the structural connectomes defined by tractography. Structural connectomes derived by covariation of cortical morphology was investigated, however, no statistically significant difference at longitudinal level was detected. Despite the belief that covariation of cortical morphology is related to anatomical connectivity of white matter^34^, the technique was not able to detect longitudinal differences in the interval of observation, most likely because these differences are more visible in the “within-brain” connectivity given by tractography, as previously suggested^19^. Therefore, these features cannot be used to perform a GWAS focused on longitudinal changes. Nevertheless, previous GWAS focused on VBM features at one time point^23^ found an association with the *APOE* gene and other SNPs related to the ephrin receptor, which are known to be correlated with the loci descried below.

In this data, Louvain modularity analysis identified the SNP rs144596626 (p-value=2.68e-10), in the *CDH18* locus, as the most significant SNP to manipulating changes in brain segregation (See Table **??** and Table 3). The *CDH18* gene encodes a cadherin that mediates calcium-dependent adhesion, playing an important role in forming the adheren junctions that bind cells. The gene is located on chromosome 5, and it is reported to be highly expressed specifically in the brain, with higher expression in different parts of the Central Nervous System (CNS), including middle temporal gyrus, cerebellum and frontal cortex^35^. The gene is associated with several neuropsychiatric disorders, as well as glioma, the most common CNV tumor among adults^36^. Looking at glioma cells, and through *in vitro* and *in vivo* functional experiments, Bai et al.^36^ showed that *CDH18* acts as a tumor suppressor through the downstream gene target *UQCRC2*, and suggested targeting *CDH18* in glioma treatmen. Moreover, *CDH18* was reported in a meta-analysis of depression personality trait association as the nearest gene to *rs349475*^37^.

On the other hand, the change in weighted global efficiency metric over time was significantly affected by the *ANTXR2* gene in chromosome 4 (see Table 2), through the imputed SNP *rs113323321* (p-value = 4.85*e* 09) with imputation accuracy of 0.743. *ANTXR2* or ANTXR cell adhesion molecule 2 (also known as *HFS*; *ISH*; *JHF*; *CMG2*; *CMG-2*) is well-known to be involved in the development of Hyaline fibromatosis syndrome (*HFS*) through certain mutations. *HFS* is a collection of rare recessive disorders forming an abnormal growth of hyalinized fibrous tissue; it affects under-skin regions on the scalp, ears, neck, face, hands, and feet. Some studies reported that *ANTXR2* mutations manipulate the normal cell interactions with the extracellular matrix, and its deleterious mutations play an essential role in causing the allelic disorders Juvenile hyaline fibromatosis (*JHF*) and infantile systemic hyalinosis (*ISH*)^38, 39^. *ANTXR2* interacts with the *LRP6* (Low-Density Lipoprotein receptor-related protein 6) gene, which is located in chromosome 12, and is known for its genetic correlation with *APOE*. Together, their genetic variants, along with the alteration in Wnt*β* signalling, might be involved in the development of late-onset AD^25^.

**Table 2.**
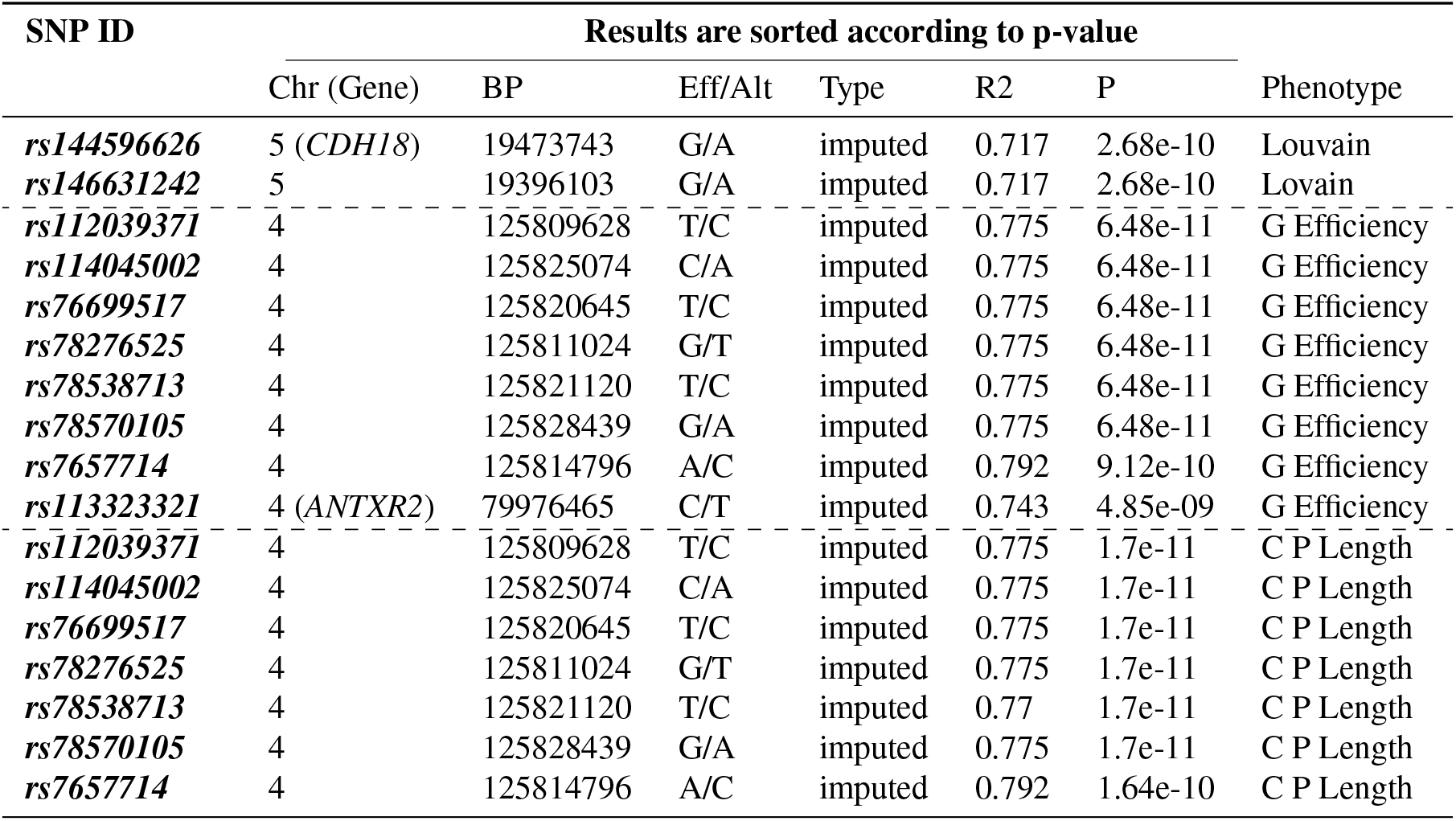
Significant associations between SNPs and global network metrics

The segregation of the brain has shown a strong relationship with the olfactory receptor (OR) family 5 (specifically, *OR5L1*, *OR5D13* and *OR5D14* - see Table 3), located in chromosome 11, through the change in brain transitivity metric. The OR act together with the odorant molecules in the nose to produce a neuronal response that recognizes smell^40^.

**Table 3.**
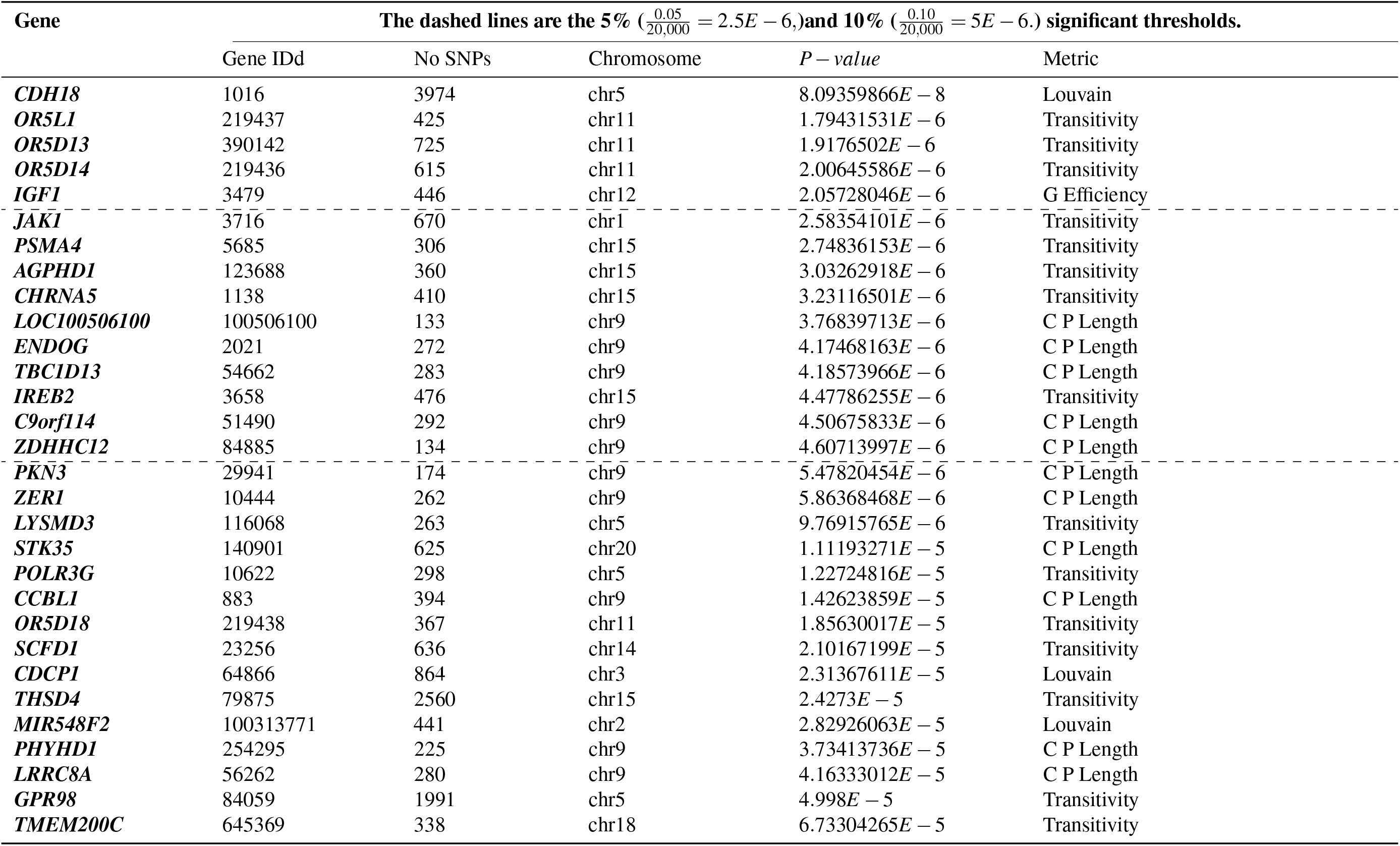
Top 30 genes: Association results with global network metrics.

Our findings also suggest that the weighted global efficiency change over time significantly associates with the insulin-like growth factor 1 (*IGF1*) gene, as shown in Table 3. A previous study in mouse brain suggests neuroprotection in a mouse model can be obtained through chronic combination therapy with EPO+IGF-I and cooperative activation of phosphatidylinositol 3-kinase/Akt/GSK-3beta signaling. However, they did not test their model in humans^26^.

At 10% significance level, we identified additional genes associated with Alzheimer’s brain segregation and integration alterations. The gene *ZDHHC12* (zinc finger DHHC-type containing 12), as with many others in chromosome 9 - including *LOC100506100*, *ENDOG*, *TBC1D13* and *C9orf114* (p-value= 3.76839713*E*–6, 4.175*E*–6, 4.186*E*–6 and 4.507*E*–6, respectively) showed a significant score (p-value=4.607E-6) in association with the change in characteristic path length (see Table 3). In an in vitro experiment,^41^ showed that *ZDHHC12* was able to alter amyloid *β*-protein precursor (*APP*) metabolism, and that the failure of AID/DHHC-12 to regulate the transportation or generation of APP in the neurons might result in the early development of AD^27^.^42^ also reported the role of Endonuclease G (*ENDOG*) in mediating the pathogenesis of neurotoxicity and striatal neuron death, through exposing the striatal neurons in mouse with Human immunodeficiency virus-1 (HIV-1) Tat_1*−*72_.

Located in chromosome 1, Janus kinase 1 (*Jak1*) shows a significant association with the change in transitivity metric. The same phenotype was also reported with other significant gene scores at 10% significance level (Table 3) such as the proteasome subunit alpha 4 (*PSMA4*) on chromosome 15, *AGPHD1*, *CHRNA5* and *IREB2* (see Fig 6). The dysregulation of the inter-cellular JAK-STAT signaling pathway, which activates *Jak1* and the Janus kinase protein family, is at the core of neurodegenerative diseases and other brain disorders^28^. *JAK2*/*STAT3* activation, in particular, was illustrated to protect the neuron, while alteration of the same pathway might play a role in developing neurodegenerative diseases.

We compared our results to previously identified genetic variants in association with Alzheimer’s (specifically SNPs), all genetic variants with p-values less than 0.01, in all global network metrics, are summarised in Table 4. We retrieved the AD SNP list from Ensembl Biomart online software^43^. Our study reported rs6026398 (*β* =-0.6496, p-value=0.000814) to be the most significant SNP associated with the change in brain segregation through Louvain modularity. The threshold we set here is 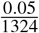, as we tested a total of 1324 pathways, though none of the SNPs passed that threshold. Our explanation for this is that variants might play a significant role in developing AD, but do not contribute that much to its progression over time. A way to take this forward is to target all genes known to affect AD susceptibility and test, in a longitudinal study design, which of them contribute to the progression of Alzheimer’s disease through the imaging features. Another recommendation is to consider studying the longitudinal association and consider whether any genetic variant has a biased contribution in different brain regions. One of the main disadvantages of this work is the sample size. We suspect the underestimation that appears in our initial GWAS results for all four phenotypes excluding transitivity, is due to sample size (Fig. **??**). In a larger sample, our result is expected to be more robust and to unveil more variants. However, to some extent, GWAS summary statistics imputation (Fig. 4 and Fig. 5) and PASCAL (6 and 7) improved this and unmasked some associations. It is worthwhile mentioning that, in this analysis, we used all the ADNI samples which satisfy our selection criteria. We also considered looking at other datasets (e.g UKBiobank and ENIGMA) but there was no data that matched our specific combination of factors required.

**Table 4.** The top 22 (p-value < 0.01) association results of AD SNPs obtained from Ensembl BioMart (no one reach the statistical threshold we set 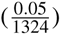.

| SNP | Results are sorted according to p-value |  |  |  |  |  |  |  |
| --- | --- | --- | --- | --- | --- | --- | --- | --- |
|  | BP | Beta | Statistic | Chr | Eff/Alt | Type(R2) | P-vale | Metric |
| *rs6026398* | 57180009 | -0.6496 | -3.544 (t) | 20 | G/A | gwas (1) | 0.000814 | Louvain |
| *rs6665019* | 25328009 | 0.834 | 3.466 (t) | 1 | A/G | gwas (1) | 0.00105 | Louvain |
| *rs2075650* | 45395619 | 0.6423 | 3.094 (t) | 19 | G/A | gwas (1) | 0.0031 | G Efficiency |
| *rs78910009* | 86408183 | NA | -2.9 (z) | 16 | G/T | imputed (0.83) | 0.00368 | C P Length |
| *rs11218343* | 121435587 | -1.025 | -3.029 (t) | 11 | C/T | gwas (1) | 0.00373 | C P Length |
| *rs4746003* | 71538292 | -0.6693 | -2.976 (t) | 10 | T/C | gwas (1) | 0.00433 | C P Length |
| *rs8014810* | 36325030 | 0.7447 | 2.975 (t) | 14 | T/G | gwas (1) | 0.00434 | Transitivity |
| *rs362389* | 73688861 | 1.284 | 2.93 (t) | 14 | C/A | gwas (1) | 0.00492 | Louvain |
| *rs73310256* | 92438849 | -1.179 | -2.916 (t) | 10 | C/T | gwas (1) | 0.00513 | G Efficiency |
| *rs4803760* | 45333834 | -0.5974 | -2.868 (t) | 19 | T/C | gwas (1) | 0.00585 | Transitivity |
| *rs11218343* | 121435587 | -0.9691 | -2.838 (t) | 11 | C/T | gwas (1) | 0.00635 | G Efficiency |
| *rs157582* | 45396219 | 0.522 | 2.836 (t) | 19 | T/C | gwas (1) | 0.00641 | G Efficiency |
| *rs362384* | 73686310 | 0.9341 | 2.834 (t) | 14 | A/C | gwas (1) | 0.00641 | Transitivity |
| *rs58920042* | 71981089 | NA | -2.72 (z) | 3 | C/T | imputed (0.729) | 0.00646 | Louvain |
| *rs117780815* | 124326227 | -1.555 | -2.793 (t) | 6 | T/A | gwas (1) | 0.0073 | Louvain |
| *rs362393* | 73689629 | 0.924 | 2.788 (t) | 14 | A/G | gwas (1) | 0.00732 | Transitivity |
| *rs1925690* | 87867063 | 0.8101 | 2.791 (t) | 6 | T/C | gwas (1) | 0.00743 | Transitivity |
| *rs7364180* | 42218856 | -0.5574 | -2.775 (t) | 22 | G/A | gwas (1) | 0.00754 | G Efficiency |
| *rs4746003* | 71538292 | -0.6278 | -2.765 (t) | 10 | T/C | gwas (1) | 0.00772 | G Efficiency |
| *rs9969729* | 108631950 | -1.04 | -2.713 (t) | 9 | A/G | gwas (1) | 0.00911 | Louvain |
| *rs117969561* | 101211189 | -1.796 | -2.702 (t) | 13 | T/C | gwas (1) | 0.00935 | C P Length |
| *rs889555* | 31122571 | -0.4928 | -2.689 (t) | 16 | T/C | gwas (1) | 0.00946 | G Efficiency |

Another concern here, is that our sample size was not sufficient to estimate the genetic correlation and heritability of our phenotypes. Most of the heritability estimation methods requires large sample sizes (at least ≈ 5k samples^44–46^) to yield robust estimates. Besides increasing the sample size, a good practice would be considering more time-points and studying the effect of genes in a survival analysis study design. In this work, we looked at the genetic variations taken at one time-point, and converted the longitudinal imaging information into a single measurement to study their association. A possible future focus would be to incorporate clinical and environmental factors such as hypertension and dementia score as well as the gene-gene and gene-environment interactions.

In summary, we conducted a longitudinal study and proposed a fast and straightforward way to quantify changes in the brain connectome through global connectivity measures of 1) segregation, through Louvain modularity and transitivity, and 2) integration. For the latter, we used two metrics including the characteristic path length and the weighted global efficiency. We conducted a genome-wide analysis, starting with four quantitative GWAS, regressing the pre-mentioned global network metric on all SNPs, and then computed the gene scores by aggregating the GWAS summary statistics at a gene-wide level. In the ADNI sample we used here, and at a power of 95%, despite the small sample size we identified significant SNPs and genes. The Louvain modularity change was affected by the *ANTXR2* gene, while through transitivity, the change in brain connectivity is associated with *OR5L1*, *OR5D13* and *OR5LD14*. On the other hand, the integration of the brain is affected by *IGF1*. A greater understanding of the genetic contribution and relationship of these genes and their effect over time through targeted studies, might facilitate the development of drug therapy to reduce the disease progression.

## Methods

### Datasets

Our analysis was conducted on the Alzheimer’s Disease Neuroimaging Initiative (ADNI) dataset publicly available at (adni.loni.usc.edu). The initiative was launched in 2003 as a public-private partnership, led by Principal Investigator Michael Weiner, MD (see www.adni-info.org for updates). To match the aim of our study, we combined two types of ADNI datasets:

1. DWI volumes was taken at two-time points, at the baseline and after 12 months (we refer to this as follow-up). In this set, we used a cohort comprised of 31 Alzheimer’s disease patients (age: 76.5 ± 7.4 years), and 49 healthy elderly subjects (77.0±5.1) matched by age, as well as 57 MCI subjects (age: 75.34 ± 5.93).
2. The PLINK binary files (BED/BIM/FAM) genotypic data for AD, controls and Early MCI.

The DWI and T1-weighted were obtained by using a GE Signa scanner 3T (General Electric, Milwaukee, WI, USA). The T1-weighted scans were acquired with voxel size = 1.2×1.0×1.0 *mm*^3^ TR = 6.984 ms; TE = 2.848 ms; flip angle=11°). DWI were acquired at voxel size = 1.4×1.4×2.7 *mm*^3^, scan time = 9 min, and 46 volumes (5 T2-weighted images with no diffusion sensitization b0 and 41 diffusion-weighted images b=1000 *s/mm*^2^).

### Preprocessing of Diffusion Imaging Data

Imaging data have T1 and DWI co-registered. To obtain the connectome; the AAL atlas^47^ is registered to the T1 volume of reference by using linear registration with 12 degrees of freedom. Despite the fact that the AAL atlas has been criticized for functional connectivity studies^48^, it has been useful in providing insights in neuroscience and physiology and it is believed to be sufficient for our case study centered on global metrics. Tractographies for all subjects were generated processing DWI data with the Python library Dipy^49^. In particular, the constant solid angle model was used^50^, and a deterministic algorithm called Euler Delta Crossings^49^ was used stemming from 2,000,000 seed-points and stopping when the fractional anisotropy was smaller than < 0.2. Tracts shorter than 30 mm, or in which a sharp angle (larger than 75°) occurred, were discarded. To construct the connectome, the graph nodes were determined using the 90 regions in the AAL atlas. Specifically, the structural connectome was built as a binary representation when more than 3 connections were given between two regions, for any pair of regions.

### Preprocessing for the Gray Matter Analysis

The data for gray matter (GM) analysis were obtained from the T1 volumes of the same subjects. The data have been preprocessed following the optimized VBM protocol from FSL^51^. Briefly, volumes have the skull stripped, bias field corrected, then are iteratively registered to a generated template in the MNI space, and have the GM segmented. During the last iteration, data are non-linearly registered to the generated template. The FSL-VBM protocol also introduces a compensation for the contraction/enlargement due to the non-linear component of the transformation: each voxel of each registered grey matter image is multiplied by the Jacobian of the warp field.

### Brain Connectivity Metrics

To assess longitudinal changes, we evaluated the following global network metrics at the two time points, at the baseline and follow-up. We then computed the absolute difference between the two measures, at each of the network metrics.

To be in line with previous work on AD and connectomics^22, 52, 53^, we focused on specific network segregation and integration features. Segregation represents the ability of a network to form communities/clusters which are well-organized^54^, while, integration represents the network’s ability to propagate information efficiently^54^.

1. Louvain modularity is a community (cluster) detection method, which iteratively transforms the network into a set of communities; each consisting of a group of nodes. Louvain modularity uses a two-step modularity optimization^17^. First, the method optimizes the modularity locally and forms communities of nodes, and secondly, it constructs a new network. The nodes of the new network are the communities formed in the previous step. These two steps are repeated iteratively until maximum modularity is obtained, and a hierarchy of communities is formed. For weighted graphs, Louvain modularity is defined as in Equation 1.

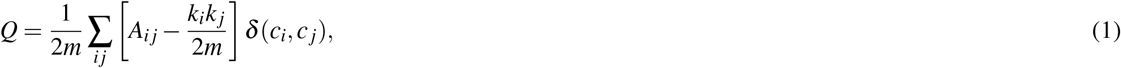

where *A_ij_* is the weight of the edge connecting nodes *i* and *j* from the adjacency matrix **A**, *k_i_* and *k_j_* are the sums of weights of the edges connected to node *i* and *j*. respectively, *m* = 1/(2*A_ij_*), *c_i_* and *c_j_* are the communities of nodes *i* and *j*, and *δ* is a simple delta function.
2. Transitivity also quantifies the segregation of a network, and is computed at a global network level as the total of all the clustering coefficients around each node in the network. It reflects the overall prevalence of clustered connectivity in a network^17^. Transitivity is mathematically defined by Equation 2.

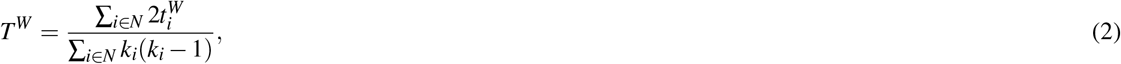

where 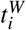 is the weighted geometric mean of triangles around node *i*, and *k_i_* its degree.
3. Weighted Global Efficiency is a network integration feature, and represents how effectively the information is exchanged over a network. This feature can be calculated as the inverse of the average weighted shortest path length in the network, as shown in Equation 3.

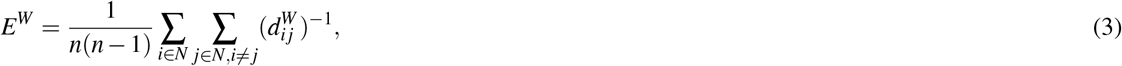

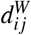, is the weighted shortest path length between node *i* and *j*, and *n* is the number of nodes.
4. Characteristic Path Length measures the integrity of the network and how fast and easily the information can flow within the network. The characteristic path length of the network is the average of all the distances between every pair in the network (see Equation 4).

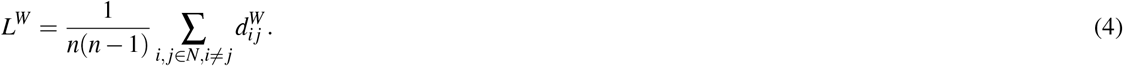

where, *d_ij_* be the number of links (connections) which represent the shortest path between node *i* and *j*. An illustrative example of global network connectivity metrics is shown in Fig. 10, the figure consists of a segregated (left) and integration (right) network.

**Figure 10.**
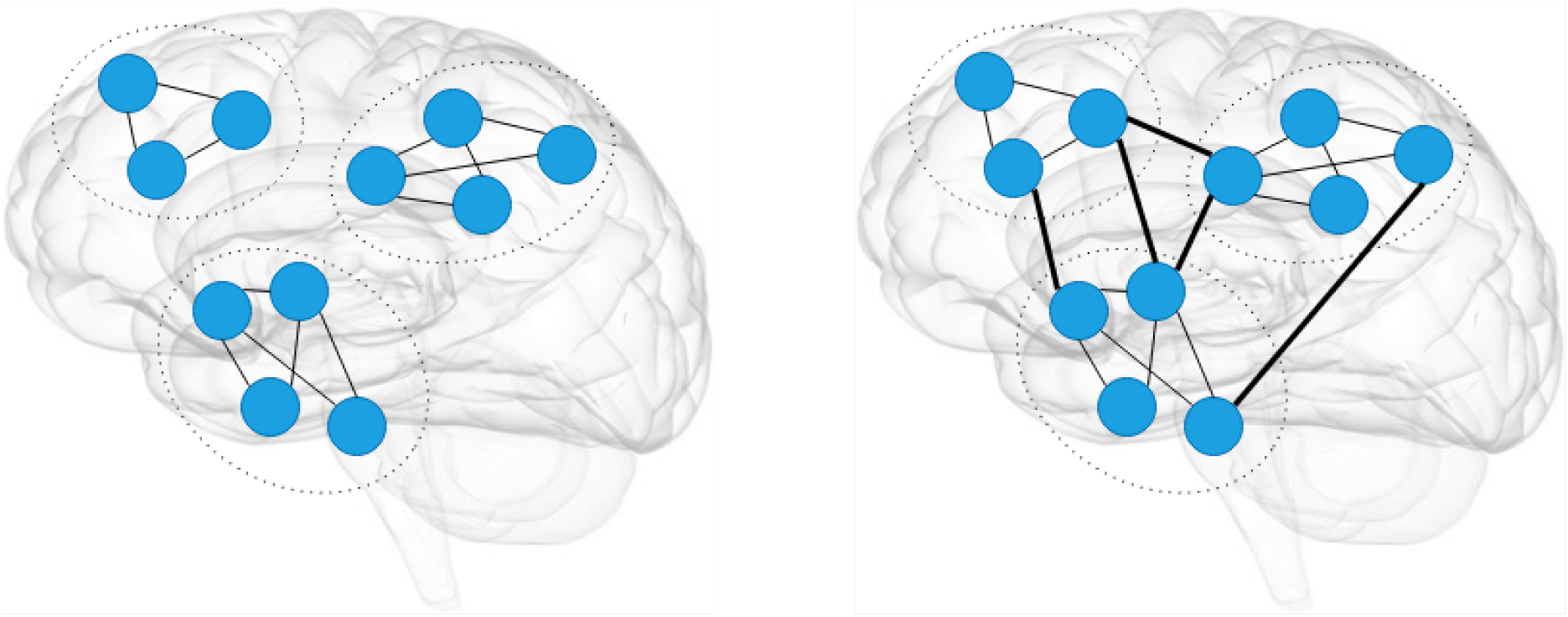
An illustrative figure of brain segregation (left) and brain integration (right). In these two figures we have the same nodes and network structure. The brain segregation represents the ability to form sub-networks as the communities on the left figures, while the integration of the brain measures the act of bringing together the different part of the brain as one connected entity, as the thick lines on the right figure.

### Gray matter analysis

Connectivity from the GM point of view is defined by the anatomical areas which covary in thickness or volume across the overall brain. Ultimately, the analysis uses another network property given by a hub index described later. Before proceeding with this analysis, a more traditional VBM pipeline was run^29^: The method, called “randomise”, performs a permutation test for the general linear model. It allows one to compare voxelwise two populations. T-statistics and corrected p-values are then computed. The comparison was carried out within the same populations (AD and control) at the two different time-points, and comparing AD against control subjects.

GM connectivity analysis follows these steps: the GM segmented and registered volumes are further subdivided into cubes of 3 × 3 × 3 voxels which now represent nodes of a network. In this way, each network has on average 6614 nodes. Edges are defined by using the Pearson correlation *r_jm_* computed between two nodes/subvolumes *v_j_* and *v_m_* each time^20^:

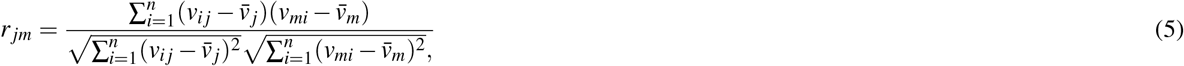

where 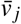, 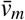 are the cubes’ mean values, and auto-correlations are set to zero. In the attempt to reduce false positives and with the aim of considering only hubs, once the connectivity matrices are constructed, these are binarized according to a threshold. We set this threshold as 2 standard deviations above the mean, though other more sophisticated threshold choices exist^19, 20^. Then, for each node, the degree of connectivity is computed by summing the binarized connections. In this way only the highly connected nodes (hubs) are defined. Lastly, values are averaged first according to the ROIs of the AAL atlas, and then for the populations at different time points. Like for the traditional VBM analysis, given the GM hubs defined at two time points we were interested in seeing whether connectivity changes occur within the interval of observations, and whether those are related to the other types of structural connectivity and gene expression.

### Integration of the two datasets

To quantify the longitudinal change in brain connectivity, we calculated the absolute difference between the baseline and follow-up for each brain connectivity metric. We then merged the absolute differences with the PLINK fam file, matching the two datasets by the subject ID.

### Quality Control

#### Quality Control: Individuals

After merging the two datasets into PLINK files, we performed some quality control procedures. First, we applied quality control at individual-level and removed all poor samples, which were identified using PLINK software and the following criteria:

1. Sex-check - here we identified all samples with ambiguous sex and removed them. We used the flag –check-sex.
2. Identifying all the individuals with missing genotype data with the flag –missing. This is to check the missingness rate of genotype information for each individual. In our data, the percentage of missingness for all individuals fell within the range (0.002834, 0.00544), since all subjects passed the threshold of 10% missingness.
3. We then identified Related Subjects (with Identity By Descent (IBD) > 20%), all subjects had IBD between 0.00 and 0.0526. We used a number of PLINK flags, including; –indep-pairwise 50 5 0.2, –extract, –genome, –min and –genome-full

After applying those quality control steps, we had a total of 57 subjects remaining for the rest of the analysis.

#### Quality Control: Genotypes

We ran quality control on the genotypes, by filtering them in terms of their minor allele frequency (MAF) with a threshold of All SNPs with less than this threshold are considered rare SNPs and were removed from the analysis. We also removed all SNPs that had missingness more than 33.33 or Genotype Call Rate < 66.67% - this was done in such a way that keeps only SNPs with sample size no less than 38. In addition, SNPs which deviate from the Hardy-Weinberg Equilibrium (HWE) were removed, these are SNPs that have p-values of less than 5e-7 in the HWE test^55^ (in total 351 SNPs did not satisfy the HWE). We used the flags, –maf, –geno and –hwe, and in total 7111195 SNPs remained.

#### Quality Control Correcting for Population Stratification

In this quality control step, we checked for the multiple presence of subpopulations in our sample. This is to make sure if we find significant variants, that the differences in allele frequencies is due to the trait under study and controls for the different ethnic groups. Population stratification helps to avoid false positives^56^. Using multiple ancestry reference genotypic information, we compared the genotypes of each study sample and estimated its ancestry with the Multi-Dimensional Scaling (MDS) analysis^57^. We observed that most of our samples belong to the Caucasian population (CEU) and therefore, proceeded by only selecting the Caucasian samples in our study. In Fig. 3, we show the genotypes of our samples compared with the reference data after the population stratification correction. We included all 57 samples as all belong to the CEU (Caucasian) ancestry. All previous quality control procedures used here followed the ENIGMA protocol^57^. The genetic reference population used here contains 13,479,643 variants that were observed more than once in the European population. These reference data were obtained by ENIGMA from the 1KGP reference set (phase 1 release v3), and imputed.

#### Quantile Normalisation of Phenotypes

Supplemenrary Fig. **??** and Supplementary Fig. **??** indicate that our phenotypes are not symmetrically distributed, and there are potential outliers. Linear models assume asymmetric distribution of the response variable. Therefore, to allow the use of linear models and conduct quantitative GWAS for our traits, we first had to normalize our phenotypes. Here, we used PLINK2^58^ (www.cog-genomics.org/plink/2.0/) to perform a quantile normalization^59^ on our phenotypes, using the flag –quantile-normalized.

### Integrated Data Analysis

#### Genome-Wide Association Analysis

We performed four quantitative GWAS separately using PLINK software^31^ (http://pngu.mgh.harvard.edu/purcell/plink/). A GWAS for each network connectivity metric measured as the absolute difference between the baseline and follow-up was performed with 57 individuals, and a total of 7111195 SNPs.

#### Statistical Thresholds

To correct for multiple testing in this analysis, and unless otherwise stated, we rely on the Bonferroni correction^60, 61^, using the simple equation below,

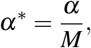

where, *M* is the number of tests, and *α* is the desired significance level. The p-values are then compared with the threshold *α*\*.

#### Imputation of GWAS Results

More quality control was done before the imputation of GWAS summary statistics using the Functionally-informed Z-score Imputation (FIZI) python tool (https://pypi.org/project/pyfizi/, https://github.com/bogdanlab/fizi). Using the munge function, 4763 SNPs with duplicated rs numbers and 85757 SNPs with N < 38.0 were removed with a remaining number of 6792416 SNPs. We then imputed the summary statistics with ImpG-Summary - Imputation from summary statistics algorithm^62^. In this step, we relied on the European 1000 Genomes^63^ haplotypes as a reference panel and performed the Gaussian imputation with FIZI. We managed to impute an additional 2222623 SNPs.

#### Gene-Wide Scores and Pathway Analysis

After we obtained the GWAS association results, we used them as input for the PASCAL software^24^ to aggregate SNPs at a gene level, and hence, compute gene scores for the four network measures. Along with the obtained association statistics PASCAL uses a reference population from the 1000 Genomes Project to correct for linkage disequilibrium (LD) between SNPs. We set PASCAL to compute the gene score as well as the pathway scores, according to the max of chi-square statistics. We got a p-value for each gene, and for each gene set (or, pathway) provided that there were SNPs presents for that gene. Finally, we used Python to plot the Manhattan plot, and R studio^64^ to plot the qq-plots. All steps are summarized in the pipeline shown in Figure 1.

## Acknowledgements

We would like to acknowledge our funders, the Organisation for Women in Science for the Developing World (OWSD), the Swedish International Development Cooperation Agency (Sida) and the University of Cape Town for their continuous support. Most of the computations in this work were performed using facilities provided by the University of Cape Town’s ICTS High Performance Computing team: http://hpc.uct.ac.za. We would like to take this opportunity to thank Dr Eleanna Kara for her valuable feedback. Data collection and sharing for this project was funded by the Alzheimer’s Disease Neuroimaging Initiative (ADNI) (National Institutes of Health Grant U01 AG024904) and DOD ADNI (Department of Defense award number W81XWH-12-2-0012). ADNI is funded by the National Institute on Aging, the National Institute of Biomedical Imaging and Bioengineering, and through generous contributions from the following: AbbVie, Alzheimer’s Association; Alzheimer’s Drug Discovery Foundation; Araclon Biotech; BioClinica, Inc.; Biogen; Bristol-Myers Squibb Company; CereSpir, Inc.; Cogstate; Eisai Inc.; Elan Pharmaceuticals, Inc.; Eli Lilly and Company; EuroImmun; F. Hoffmann-La Roche Ltd and its affiliated company Genentech, Inc.; Fujirebio; GE Healthcare; IXICO Ltd.;Janssen Alzheimer Immunotherapy Research & Development, LLC.; Johnson & Johnson Pharmaceutical Research & Development LLC.; Lumosity; Lundbeck; Merck & Co., Inc.;Meso Scale Diagnostics, LLC.; NeuroRx Research; Neurotrack Technologies; Novartis Pharmaceuticals Corporation; Pfizer Inc.; Piramal Imaging; Servier; Takeda Pharmaceutical Company; and Transition Therapeutics. The Canadian Institutes of Health Research is providing funds to support ADNI clinical sites in Canada. Private sector contributions are facilitated by the Foundation for the National Institutes of Health (http://www.fnih.org). The grantee organization is the Northern California Institute for Research and Education, and the study is coordinated by the Alzheimer’s Therapeutic Research Institute at the University of Southern California. ADNI data are disseminated by the Laboratory for Neuro Imaging at the University of Southern California.

## Author Contributions Statement

A.C. ran all neuroimaging analysis, S.S.M.E. conducted the imaging genetic analysis and wrote the paper with A.C., N.J.M. and A.C gave continued feedback and closely followed-up the progress. E.R.C. gave constructive feedback on the analysis, and all authors proofread the final version of the paper.

## Additional Information

### Data and Code Availability

The data used in this work is available at the ADNI repository (http://adni.loni.usc.edu/).

The used code is publicly available at the url https://github.com/elssam/GWAS-BCCAD.

### Supplementary information

Supplementary materials available for this paper.

## References

1. Raj, A. et al. Network diffusion model of progression predicts longitudinal patterns of atrophy and metabolism in Alzheimer’s disease. Cell reports 10, 359–369 (2015).

2. Dominy, S. S. et al. Porphyromonas gingivalis in alzheimer’s disease brains: Evidence for disease causation and treatment with small-molecule inhibitors. Sci. advances 5, eaau3333 (2019).

3. Ballard, C. et al. Alzheimer’s disease. The Lancet 377, 1019 – 1031 (2011).

4. Corder, E., Saunders, A. et al. Gene dose of apolipoprotein E type 4 allele and the risk of Alzheimer’s disease in late onset families. Science 261, 921–923 (1993).

5. Lambert, J.-C. et al. Meta-analysis of 74,046 individuals identifies 11 new susceptibility loci for alzheimer’s disease. Nat. genetics 45, 1452 (2013).

6. Bertram, L. & other. The genetics of Alzheimer disease: back to the future. Neuron 68, 270–281 (2010).

7. Li, H. et al. Candidate single-nucleotide polymorphisms from a genomewide association study of alzheimer disease. Arch. neurology 65, 45–53 (2008).

8. Naj, A. C. et al. Dementia revealed: novel chromosome 6 locus for late-onset alzheimer disease provides genetic evidence for folate-pathway abnormalities. PLoS genetics 6, e1001130 (2010).

9. Mier, W. & Mier, D. Advantages in functional imaging of the brain. Front. human neuroscience 9, 249 (2015).

10. Thompson, P. M., Hibar, D. P., Stein, J. L., Prasad, G. & Jahanshad, N. Genetics of the connectome and the enigma project. In Micro-, Meso-and Macro-Connectomics of the Brain, 147–164 (Springer, 2016).

11. Stein, J. L. et al. Voxelwise genome-wide association study (vgwas). neuroimage 53, 1160–1174 (2010).

12. Jack Jr, C. R. et al. The alzheimer’s disease neuroimaging initiative (adni): Mri methods. J. Magn. Reson. Imaging: An Off. J. Int. Soc. for Magn. Reson. Medicine 27, 685–691 (2008).

13. Stein, J. L. et al. Identification of common variants associated with human hippocampal and intracranial volumes. Nat. genetics 44, 552 (2012).

14. Hibar, D. P. et al. Common genetic variants influence human subcortical brain structures. Nature 520, 224 (2015).

15. Fang, J. et al. Joint sparse canonical correlation analysis for detecting differential imaging genetics modules. Bioinformatics 32, 3480–3488 (2016).

16. Hagmann, P. et al. Mapping the structural core of human cerebral cortex. PLoS biology 6, e159 (2008).

17. Rubinov, M. & Sporns, O. Complex network measures of brain connectivity: uses and interpretations. Neuroimage 52, 1059–1069 (2010).

18. Good, C. D. et al. A voxel-based morphometric study of ageing in 465 normal adult human brains. Neuroimage 14, 21–36 (2001).

19. Forsberg, L., Sigurdsson, S., Launer, L. J., Gudnason, V. & Ullén, F. Structural covariability hubs in old age. NeuroImage 189, 307–315 (2019).

20. Tijms, B. M., Seriès, P., Willshaw, D. J. & Lawrie, S. M. Similarity-based extraction of individual networks from gray matter mri scans. Cereb. cortex 22, 1530–1541 (2012).

21. Wu, K. et al. A longitudinal study of structural brain network changes with normal aging. Front. human neuroscience 7, 113 (2013).

22. Jahanshad, N. et al. Genome-wide scan of healthy human connectome discovers spon1 gene variant influencing dementia severity. Proc. Natl. Acad. Sci. 110, 4768–4773 (2013).

23. Shen, L. et al. Whole genome association study of brain-wide imaging phenotypes for identifying quantitative trait loci in mci and ad: A study of the adni cohort. Neuroimage 53, 1051–1063 (2010).

24. Lamparter, D., Marbach, D., Rueedi, R., Kutalik, Z. & Bergmann, S. Fast and rigorous computation of gene and pathway scores from snp-based summary statistics. PLoS computational biology 12, e1004714 (2016).

25. De Ferrari, G. V. et al. Common genetic variation within the low-density lipoprotein receptor-related protein 6 and late-onset alzheimer’s disease. Proc. Natl. Acad. Sci. 104, 9434–9439 (2007).

26. Kang, Y.-J. et al. Erythropoietin plus insulin-like growth factor-i protects against neuronal damage in a murine model of human immunodeficiency virus-associated neurocognitive disorders. Annals neurology 68, 342–352 (2010).

27. Young, F. B., Butland, S. L., Sanders, S. S., Sutton, L. M. & Hayden, M. R. Putting proteins in their place: palmitoylation in huntington disease and other neuropsychiatric diseases. Prog. neurobiology 97, 220–238 (2012).

28. Nicolas, C. S. et al. The role of jak-stat signaling within the cns. Jak-Stat 2, e22925 (2013).

29. Winkler, A. M., Ridgway, G. R., Webster, M. A., Smith, S. M. & Nichols, T. E. Permutation inference for the general linear model. Neuroimage 92, 381–397 (2014).

30. Wang, X.-H., Jiao, Y. & Li, L. Mapping individual voxel-wise morphological connectivity using wavelet transform of voxel-based morphology. PloS one 13, e0201243 (2018).

31. Purcell, S. et al. Plink: a tool set for whole-genome association and population-based linkage analyses. The Am. J. Hum. Genet. 81, 559–575 (2007).

32. Cuyvers, E. & Sleegers, K. Genetic variations underlying alzheimer’s disease: evidence from genome-wide association studies and beyond. The Lancet Neurol. 15, 857–868 (2016).

33. Elsheikh, S., Chimusa, E. R., Mulder, N. & Crimi, A. Relating connectivity changes in brain networks to genetic information in alzheimer patients. In 2018 IEEE 15th International Symposium on Biomedical Imaging (ISBI 2018), 1390–1393 (IEEE, 2018).

34. Pezawas, L. et al. The brain-derived neurotrophic factor val66met polymorphism and variation in human cortical morphology. J. Neurosci. 24, 10099–10102 (2004).

35. Fagerberg, L. et al. Analysis of the human tissue-specific expression by genome-wide integration of transcriptomics and antibody-based proteomics. Mol. & Cell. Proteomics 13, 397–406 (2014).

36. Bai, Y.-h. et al. A novel tumor-suppressor, cdh18, inhibits glioma cell invasiveness via uqcrc2 and correlates with the prognosis of glioma patients. Cell. Physiol. Biochem. 48, 1755–1770 (2018).

37. Terracciano, A. et al. Genome-wide association scan of trait depression. Biol. psychiatry 68, 811–817 (2010).

38. Dowling, O. et al. Mutations in capillary morphogenesis gene-2 result in the allelic disorders juvenile hyaline fibromatosis and infantile systemic hyalinosis. The Am. J. Hum. Genet. 73, 957–966 (2003).

39. Hanks, S. et al. Mutations in the gene encoding capillary morphogenesis protein 2 cause juvenile hyaline fibromatosis and infantile systemic hyalinosis. The Am. J. Hum. Genet. 73, 791–800 (2003).

40. O’Leary, N. A. et al. Reference sequence (refseq) database at ncbi: current status, taxonomic expansion, and functional annotation. Nucleic acids research 44, D733–D745 (2015).

41. Mizumaru, C. et al. Suppression of app-containing vesicle trafficking and production of *β*-amyloid by aid/dhhc-12 protein. J. neurochemistry 111, 1213–1224 (2009).

42. Singh, I. N. et al. Apoptotic death of striatal neurons induced by human immunodeficiency virus-1 tat and gp120: Differential involvement of caspase-3 and endonuclease g. J. neurovirology 10, 141–151 (2004).

43. Kinsella, R. J. et al. Ensembl biomarts: a hub for data retrieval across taxonomic space. Database 2011 (2011).

44. Bulik-Sullivan, B. et al. An atlas of genetic correlations across human diseases and traits. Nat. genetics 47, 1236 (2015).

45. Finucane, H. K. et al. Partitioning heritability by functional annotation using genome-wide association summary statistics. Nat. genetics 47, 1228 (2015).

46. Yang, J. et al. Common snps explain a large proportion of the heritability for human height. Nat. genetics 42, 565 (2010).

47. Tzourio-Mazoyer, N. et al. Automated anatomical labeling of activations in spm using a macroscopic anatomical parcellation of the mni mri single-subject brain. Neuroimage 15, 273–289 (2002).

48. Gordon, E. M. et al. Generation and evaluation of a cortical area parcellation from resting-state correlations. Cereb. cortex 26, 288–303 (2014).

49. Garyfallidis, E. et al. Dipy, a library for the analysis of diffusion mri data. Front. Neuroinformatics 8, DOI: 10.3389/fninf.2014.00008 (2014).

50. Aganj, I. et al. Reconstruction of the orientation distribution function in single-and multiple-shell q-ball imaging within constant solid angle. Magn. resonance medicine 64, 554–566 (2010).

51. Douaud, G. et al. Anatomically related grey and white matter abnormalities in adolescent-onset schizophrenia. Brain 130, 2375–2386 (2007).

52. Prasad, G., Nir, T., Toga, A. & Thompson, P. Tractography density and network measures in Alzheimer’s disease. In Biomedical Imaging, 2013 IEEE 10th International Symposium on, 692–695 (2013).

53. Brown, J. et al. Bain network local interconnectivity loss in aging apoe-4 allele carriers. Proc. Natl. Acad. Sci. 108, 20760–20765 (2011).

54. Deco, G., Tononi, G. et al. Rethinking segregation and integration: contributions of whole-brain modelling. Nat. Rev. Neurosci. 16, 430–439 (2015).

55. Wigginton, J. E., Cutler, D. J. & Abecasis, G. R. A note on exact tests of hardy-weinberg equilibrium. The Am. J. Hum. Genet. 76, 887–893 (2005).

56. Hamer, D. & Sirota, L. Beware the chopsticks gene (2000).

57. Egs, T. Enigma2 1kgp cookbook (v3). Enhancing Neuroimaging Genet. through MetaAnalysis (ENIGMA) Consortium (2013).

58. Chang, C. C. et al. Second-generation plink: rising to the challenge of larger and richer datasets. Gigascience 4, 7 (2015).

59. Bolstad, B. M., Irizarry, R. A., Åstrand, M. & Speed, T. P. A comparison of normalization methods for high density oligonucleotide array data based on variance and bias. Bioinformatics 19, 185–193 (2003).

60. White, T., van der Ende, J. & Nichols, T. E. Beyond bonferroni revisited: concerns over inflated false positive research findings in the fields of conservation genetics, biology, and medicine. Conserv. Genet. 1–11 (2019).

61. Narum, S. R. Beyond bonferroni: less conservative analyses for conservation genetics. Conserv. genetics 7, 783–787 (2006).

62. Pasaniuc, B. et al. Fast and accurate imputation of summary statistics enhances evidence of functional enrichment. Bioinformatics 30, 2906–2914 (2014).

63. Consortium,. G. P. et al. An integrated map of genetic variation from 1,092 human genomes. Nature 491, 56 (2012).

64. RStudio Team. RStudio: Integrated Development Environment for R. RStudio, Inc., Boston, MA (2015).

