## Supplemental_Figures_and_Tables for "Genome-Wide Association Study of Brain Connectivity Changes for Alzheimer’s Disease"

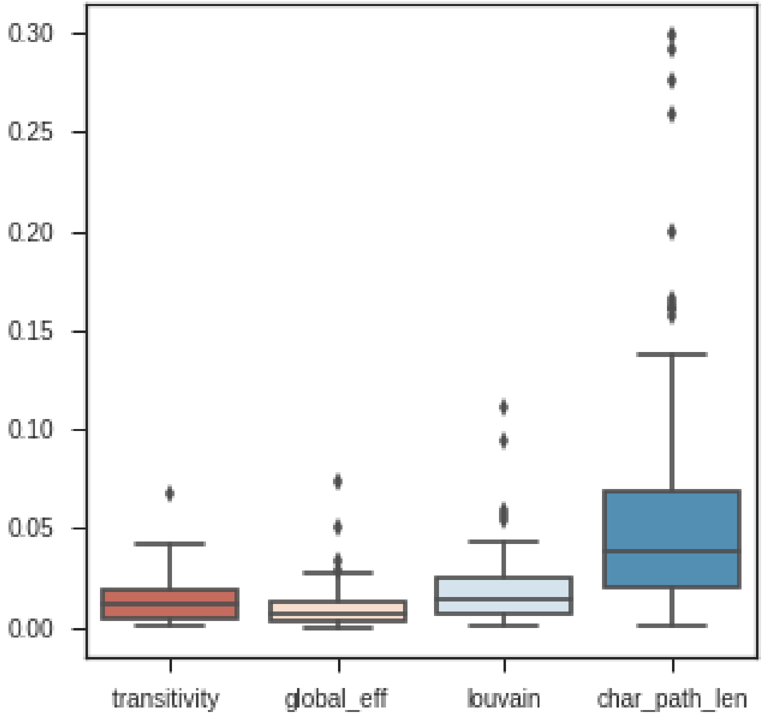


**Figure S1.** Distribution of global network metrics for controls, MCI and AD subjects, combined. Shortcuts stand for; Louvain: Louvain modularity, global_eff: global effeciency, and char_path_len: characteristic path length

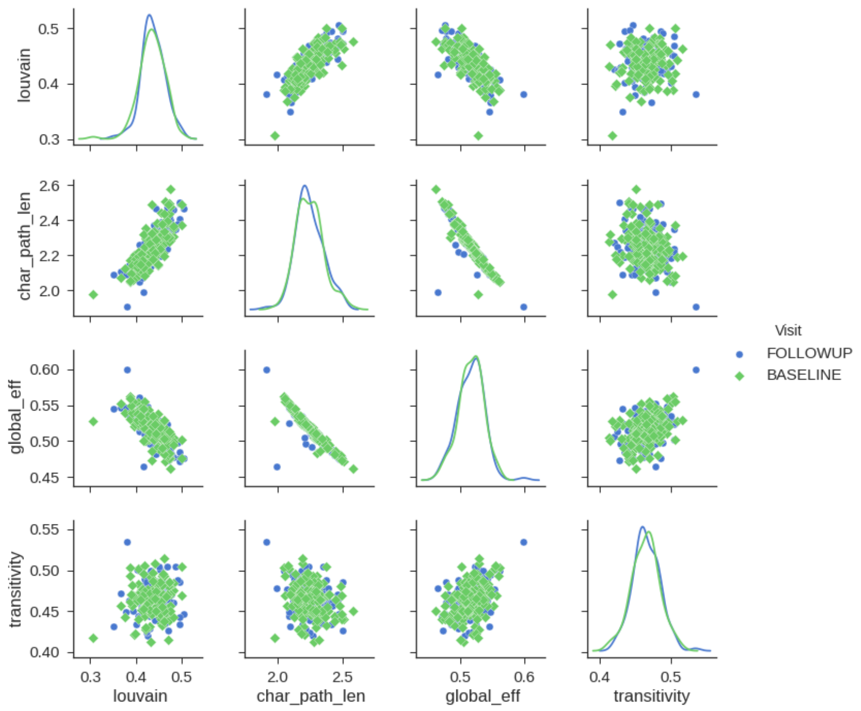


**Figure S2.** Global network metrics scatter plots: The sub-plots compare the four global network metrics before and after 12 months (baseline vs follow-up). Diagonal plots show the distribution of the actual metrics in the baseline and follow-up, while the remaining plots show the correlation between the metrics, for all participants.

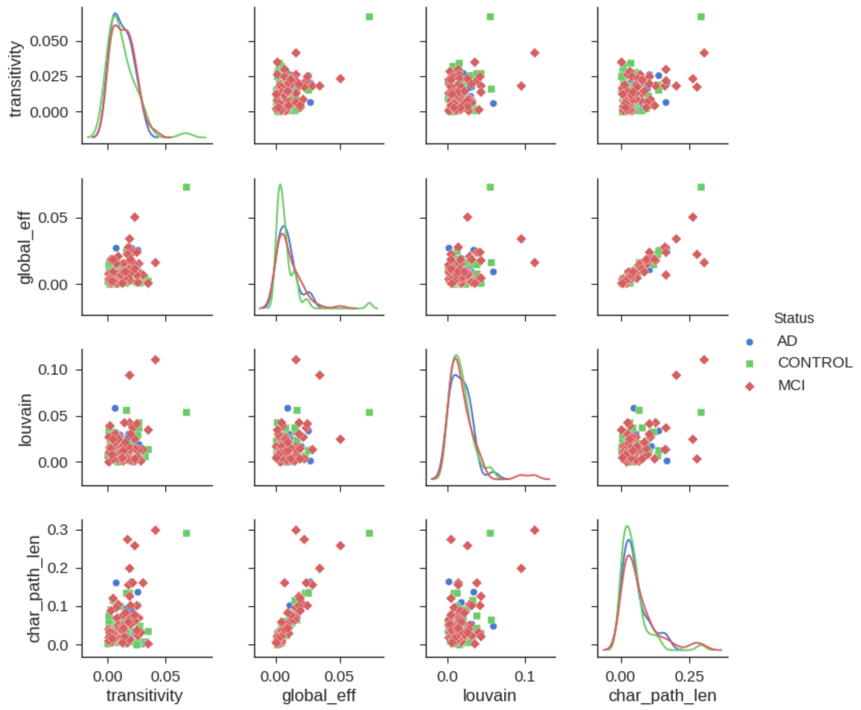


**Figure S3.** Global network metrics scatter plots: The sub-figures show the distribution of the absolute difference of the four network metrics (diagonal plots); as well as the pairwise correlation between them (remaining plots). Each plot compares AD, MCI and controls.

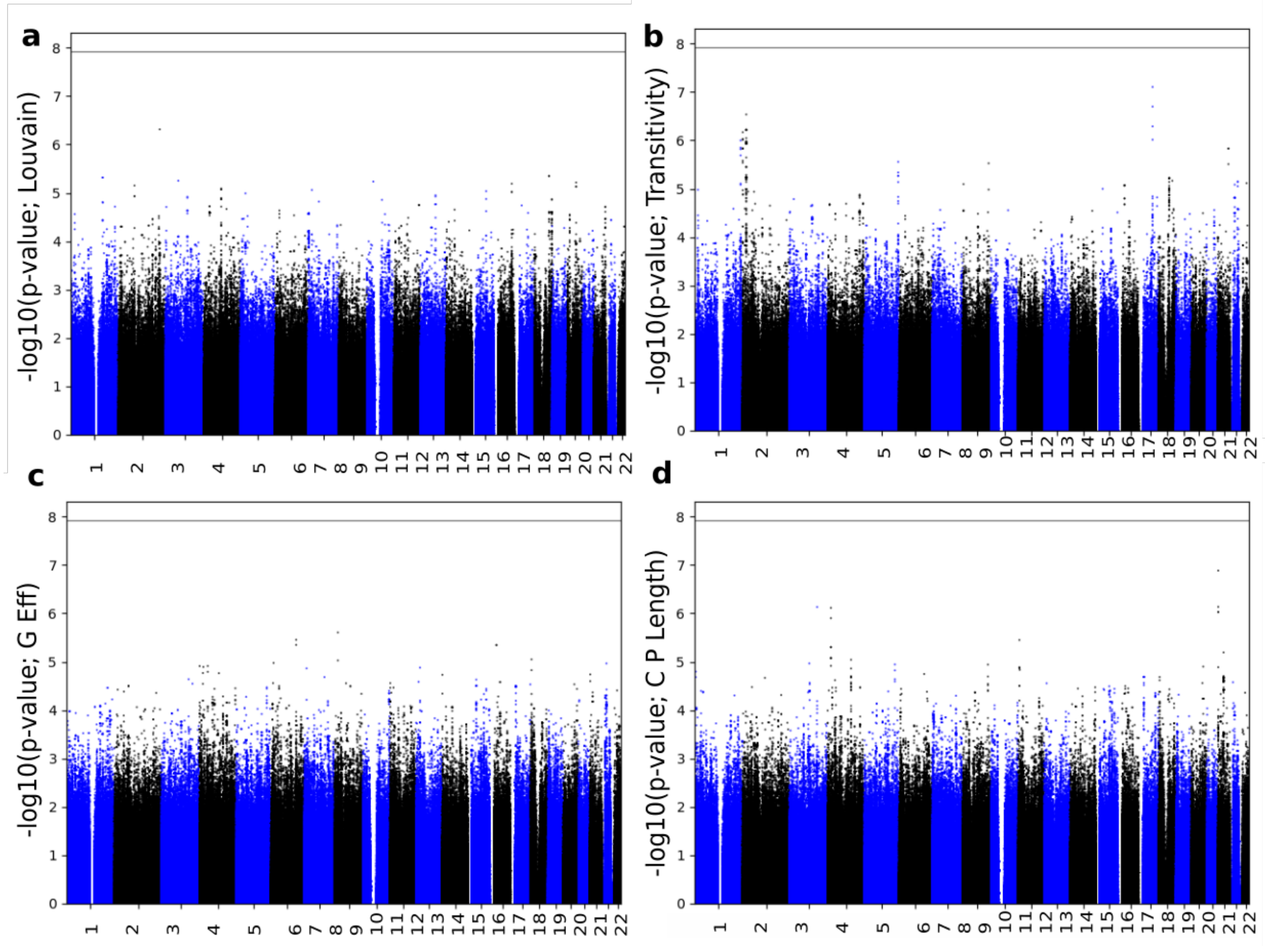


**Figure S4.** Manhattan plots of GWAS results for the change in Louvain modularity (a) and transitivity (b) global efficiency (c) and the change in characteristics path length (d) integration and segregation connectivity metrics.

**Table S1.** Louvain modularity GWAS results: Top 15 SNPs

| SNP | Dashed line is the 5% threshold | | | | | | |
|---|---|---|---|---|---|---|---|
| | Chr | BP | Eff/Alt | Type(R2) | Statistic | $\beta$ | P |
| ***rs144596626*** | 5 | 19473852 | G/A | imputed (0.717) | 6.32 (z) | | 2.68e-10 |
| ***rs146631242*** | 5 | 19396212 | G/A | imputed (0.717) | 6.32 (z) | | 2.68e-10 |
| ***rs35942723*** | 2 | 67943399 | T/C | imputed (0.7) | 5.43 (z) | | 5.5e-08 |
| ***rs2460661*** | 5 | 98903041 | C/T | imputed (0.65) | 5.3 (z) | | 1.14e-07 |
| ***rs144454897*** | 3 | 45220242 | G/A | imputed (0.839) | -5.2 (z) | | 2.03e-07 |
| ***rs12694279*** | 2 | 213253406 | C/A | gwas (1) | -5.992 (t) | -1.19 | 4.85e-07 |
| ***rs146293495*** | 11 | 87983529 | C/T | imputed (0.782) | -5.01 (z) | | 5.54e-07 |
| ***rs145955468*** | 4 | 182593928 | G/A | imputed (0.818) | 4.88 (z) | | 1.07e-06 |
| ***rs185097390*** | 4 | 182653779 | G/A | imputed (0.818) | 4.88 (z) | | 1.07e-06 |
| ***rs149021889*** | 5 | 19168938 | T/G | imputed (0.603) | 4.82 (z) | | 1.42e-06 |
| ***rs189745822*** | 7 | 130949241 | C/A | imputed (0.805) | -4.8 (z) | | 1.56e-06 |
| ***rs193071172*** | 7 | 130946162 | G/A | imputed (0.805) | -4.8 (z) | | 1.56e-06 |
| ***rs150749209*** | 7 | 42493974 | T/C | imputed (0.714) | -4.66 (z) | | 3.12e-06 |
| ***rs189358029*** | 17 | 43702340 | C/T | imputed (0.437) | 4.62 (z) | | 3.87e-06 |
| ***rs8053032*** | 16 | 75181579 | G/C | gwas (1) | -5.199 (t) | -0.9548 | 4.5e-06 |

**Table S2.** Transitivity GWAS results: Top 15 SNPs

| SNP | Dashed line is the 5% threshold | | | | | | P |
|---|---|---|---|---|---|---|---|
| | Chr | BP | Eff/Alt | Type(R2) | Statistic | $\beta$ | P |
| ***rs4617614*** | 11 | 55537046 | C/T | imputed (0.717) | 5.54 (z) | | 3.11e-08 |
| ***rs144573130*** | 1 | 65415375 | C/T | imputed (0.82) | 5.53 (z) | | 3.27e-08 |
| ***rs111650215*** | 15 | 78828083 | A/G | imputed (0.661) | -5.43 (z) | | 5.69e-08 |
| ***rs112671439*** | 15 | 78819074 | T/C | imputed (0.661) | -5.43 (z) | | 5.69e-08 |
| ***rs113809575*** | 15 | 78833209 | G/A | imputed (0.661) | -5.43 (z) | | 5.69e-08 |
| ***rs113882269*** | 15 | 78833286 | A/G | imputed (0.661) | -5.43 (z) | | 5.69e-08 |
| ***rs11912587*** | 22 | 38371933 | A/C | imputed (0.738) | 5.37 (z) | | 7.81e-08 |
| ***rs4459504*** | 15 | 71883930 | G/A | gwas (1) | 6.192(t) | 0.8938 | 7.83e-08 |
| ***rs144750443*** | 5 | 89847749 | C/T | imputed (0.745) | -5.2 (z) | | 1.96e-07 |
| ***rs3923493*** | 15 | 71866632 | T/C | gwas (1) | 5.973 (t) | 0.9766 | 2e-07 |
| ***rs2518679*** | 14 | 31252534 | T/C | imputed (0.679) | 5.18 (z) | | 2.21e-07 |
| ***rs61156477*** | 14 | 31266735 | T/C | imputed (0.679) | 5.18 (z) | | 2.21e-07 |
| ***rs77762911*** | 14 | 31255027 | T/G | imputed (0.679) | 5.18 (z) | | 2.21e-07 |
| ***rs8018229*** | 14 | 31261372 | G/A | imputed (0.679) | 5.18 (z) | | 2.21e-07 |
| ***rs147801202*** | 2 | 19102920 | A/ATG | gwas (1) | -5.939 (t) | -1.049 | 2.91e-07 |

**Table S3.** Global efficiency GWAS results: Top 15 SNPs

| SNP | Dashed line is the 5% threshold | | | | | | P |
|---|---|---|---|---|---|---|---|
| | Chr | BP | Eff/Alt | Type(R2) | Statistic | $\beta$ | P |
| ***rs112039371*** | 4 | 126730783 | T/C | imputed (0.775) | -6.53 (z) | | 6.48e-11 |
| ***rs114045002*** | 4 | 126746229 | C/A | imputed (0.775) | -6.53 (z) | | 6.48e-11 |
| ***rs76699517*** | 4 | 126741800 | T/C | imputed (0.775) | -6.53 (z) | | 6.48e-11 |
| ***rs78276525*** | 4 | 126732179 | G/T | imputed (0.775) | -6.53 (z) | | 6.48e-11 |
| ***rs78538713*** | 4 | 126742275 | T/C | imputed (0.775) | -6.53 (z) | | 6.48e-11 |
| ***rs78570105*** | 4 | 126749594 | G/A | imputed (0.775) | -6.53 (z) | | 6.48e-11 |
| ***rs7657714*** | 4 | 126735951 | A/C | imputed (0.792) | -6.12 (z) | | 9.12e-10 |
| ***rs113323321*** | 4 | 80897619 | C/T | imputed (0.743) | 5.85 (z) | | 4.85e-09 |
| ***rs192963808*** | 12 | 102764731 | A/G | imputed (0.694) | 5.53 (z) | | 3.28e-08 |
| ***rs146655189*** | 12 | 102674455 | C/T | imputed (0.717) | 5.48 (z) | | 4.27e-08 |
| ***rs148061827*** | 10 | 109846291 | G/T | imputed (0.678) | 4.84 (z) | | 1.28e-06 |
| ***rs2139572*** | 12 | 102767660 | C/T | imputed (0.657) | 4.82 (z) | | 1.44e-06 |
| ***rs149903755*** | 14 | 100808726 | A/G | imputed (0.804) | -4.77 (z) | | 1.86e-06 |
| ***rs149119261*** | 10 | 108038891 | C/T | imputed (0.762) | 4.73 (z) | | 2.3e-06 |
| ***rs62497351*** | 8 | 17086601 | G/T | gwas (1) | 5.278 (t) | 1.402 | 2.48e-06 |

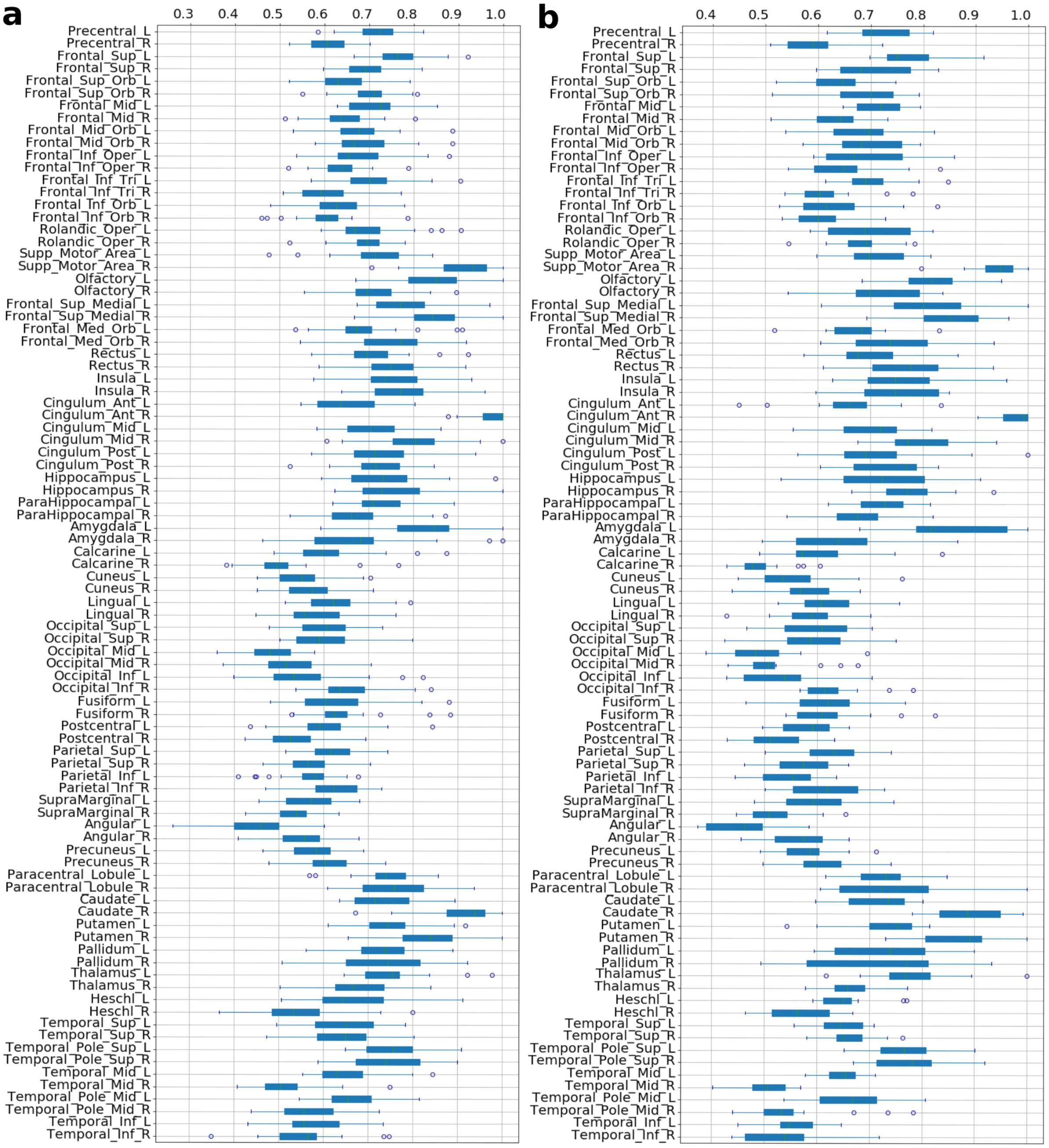


**Figure S5.** Boxplots of hubs degree centrality averaged according to the single ROI of the AAL atlas. On the left (a) are the values for the AD subjects at baseline, and (b) on the right are the values for the AD subjects at follow-up.

**Table S4.** Charactristic path length GWAS results: Top 15 SNPs

| SNP | Dashed line is the 5% threshold | | | | | | |
|---|---|---|---|---|---|---|---|
| | Chr | BP | Eff/Alt | Type(R2) | Statistic | $\beta$ | P |
| ***rs112039371*** | 4 | 126730783 | T/C | imputed (0.775) | -6.73 (z) | | 1.7e-11 |
| ***rs114045002*** | 4 | 126746229 | C/A | imputed (0.775) | -6.73 (z) | | 1.7e-11 |
| ***rs76699517*** | 4 | 126741800 | T/C | imputed (0.775) | -6.73 (z) | | 1.7e-11 |
| ***rs78276525*** | 4 | 126732179 | G/T | imputed (0.775) | -6.73 (z) | | 1.7e-11 |
| ***rs78538713*** | 4 | 126742275 | T/C | imputed (0.775) | -6.73 (z) | | 1.7e-11 |
| ***rs78570105*** | 4 | 126749594 | G/A | imputed (0.775) | -6.73 (z) | | 1.7e-11 |
| ***rs7657714*** | 4 | 126735951 | A/C | imputed (0.792) | -6.39 (z) | | 1.64e-10 |
| ***rs10113946*** | 9 | 131534333 | C/T | imputed (0.725) | -5.36 (z) | | 8.29e-08 |
| ***rs11560592*** | 9 | 131532694 | C/T | imputed (0.725) | -5.36 (z) | | 8.29e-08 |
| ***rs28521006*** | 9 | 131534909 | G/A | imputed (0.725) | -5.36 (z) | | 8.29e-08 |
| ***rs35354551*** | 20 | 2112390 | C/T | gwas (1) | -6.089 (t) | -1.107 | 1.31e-07 |
| ***rs113323321*** | 4 | 80897619 | C/T | imputed (0.743) | 5.22 (z) | | 1.83e-07 |
| ***rs10122433*** | 9 | 131577388 | A/C | imputed (0.753) | -5.18 (z) | | 2.25e-07 |
| ***rs12236573*** | 9 | 131555366 | G/A | imputed (0.715) | -5.03 (z) | | 5.01e-07 |
| ***rs10115869*** | 9 | 131652502 | G/A | imputed (0.735) | -5.03 (z) | | 5.03e-07 |

**Table S5.** Top 20 gene sets (pathways) results derived from GWAS summary statistics of global network metrics.

| **Gene Set** | **Results are sorted by p-value.** | | |
|---|---|---|---|
| | Chr | Pvalue | Metric |
| **REACTOME_BIOLOGICAL_OXIDATIONS** | 10 | 2.91E-5 | Louvain |
| **REACTOME_REGULATION_OF_ORNITHINE_DECARBOXYLASE_ODC** | 15 | 5.2E-5 | Transitivity |
| **KEGG_ALDOSTERONE_REGULATED_SODIUM_REABSORPTION** | 12 | 6.6E-5 | G Efficiency |
| **BIOCARTA_IL7_PATHWAY** | 1 | 6.9E-5 | Transitivity |
| **REACTOME_IL_6_SIGNALING** | 1 | 7.6E-5 | Transitivity |
| **KEGG_PATHWAYS_IN_CANCER** | 12 | 1.05E-4 | G Efficiency |
| **BIOCARTA_ERYTH_PATHWAY** | 12 | 1.32E-4 | G Efficiency |
| **BIOCARTA_BAD_PATHWAY** | 12 | 1.38E-4 | G Efficiency |
| **KEGG_PROGESTERONE_MEDIATED_OOCYTE_MATURATION** | 12 | 1.38E-4 | G Efficiency |
| **BIOCARTA_IL22BP_PATHWAY** | 1 | 1.57E-4 | Transitivity |
| **REACTOME_GENERIC_TRANSCRIPTION_PATHWAY** | 2 | 2.03E-4 | G Efficiency |
| **KEGG_PROSTATE_CANCER** | 12 | 2.46E-4 | G Efficiency |
| **REACTOME_ACTIVATION_OF_NMDA_RECEPTOR_ UPON_GLUTAMATE_BINDING_AND_POSTSYNAPTIC_EVENTS** | 8 | 2.56E-4 | Transitivity |
| **REACTOME_POST_NMDA_RECEPTOR_ACTIVATION_EVENTS** | 8 | 2.7E-4 | Transitivity |
| **KEGG_LYSOSOME** | 11 | 3.8E-4 | Louvain |
| **REACTOME_OLFACTORY_SIGNALING_PATHWAY** | 11 | 4.08E-4 | Transitivity |
| **REACTOME_ANTIVIRAL_MECHANISM_BY_IFN_STIMULATED_GENES** | 1 | 4.17E-4 | Transitivity |
| **REACTOME_REGULATION_OF_IFNA_SIGNALING** | 1 | 4.27E-4 | Transitivity |
| **BIOCARTA_IL2_PATHWAY** | 1 | 4.29E-4 | Transitivity |
| **KEGG_GLIOMA** | 12 | 4.34E-4 | G Efficiency |
